# pH Effect on Ligand Binding to an Enzyme Active Site

**DOI:** 10.1101/2022.07.01.498456

**Authors:** Kushal Singh, Aswathy N. Muttathukattil, Prashant Chandra Singh, Govardhan Reddy

## Abstract

Understanding the mechanism of ligands binding to their protein targets and the influence of various factors governing the binding thermodynamics is essential for rational drug design. The solution pH is one of the critical factors that can influence ligand binding to a protein cavity, especially in enzymes whose function is sensitive to the pH. Using computer simulations, we studied the pH effect on the binding of a guanidinium ion (Gdm^+^) to the active site of hen-egg white lysozyme (HEWL). HEWL serves as a model system for enzymes with two acidic residues in the active site and ligands with Gdm^+^ moieties, which can bind to the active sites of such enzymes and are present in several approved drugs treating various disorders. The computed free energy surface (FES) shows that Gdm^+^ binds to the HEWL active site using two dominant binding pathways populating multiple intermediates. We show that the residues close to the active site that can anchor the ligand could play a critical role in ligand binding. Using a Markov state model, we quantified the lifetimes and kinetic pathways connecting the different states in the FES. The protonation and deprotonation of the acidic residues in the active site in response to the pH change strongly influence the Gdm^+^ binding. There is a sharp jump in the ligand-binding rate constant when the pH approaches the largest p*K*_a_ of the acidic residue present in the active site. The simulations reveal that, at most, three Gdm^+^ can bind at the active site, with the Gdm^+^ bound in the cavity of the active site acting as a scaffold for the other two Gdm^+^ ions binding. This result implies the possibility of designing single large molecules containing multiple Gdm^+^ moieties that can have high binding affinities to inhibit the function of enzymes with two acidic residues in their active site.

## Introduction

A protein’s biological activity is mediated through its interaction with other molecules, such as polypeptides, nucleic acids, membrane lipids, ions, metabolites, etc., present in the cellular medium. These interactions can be modulated to regulate protein activity, which is exploited in drug discovery to treat disorders related to the protein. The therapeutic effect of a drug molecule is due to its binding in a specific protein site and perturbing the protein’s activity by interfering with its interaction with other molecules in the cell. Knowledge of the molecular interactions between the drug and the protein and the underlying protein-ligand binding thermodynamics and kinetics is essential for rational drug design.

External factors such as temperature, pressure, molecular crowding, pH, etc., strongly influence drug binding kinetics. In biological systems, pH plays a critical role in dictating the efficacy of a drug binding to a protein, especially enzymes. Generally, enzymes operate optimally in a specific pH range, which must be considered while designing drugs targeting them. For example, aspartic proteases such as *β*-secretase (pH = 3.5 - 5.5),^1^ HIV-1 protease (pH = 4 - 6),^2^ and influenza neuraminidases (pH = 5.5 - 6.5)^3^ operate in a narrow pH range.

As the p*K*_a_ values of the acidic residue side chains are dependent on their location in the protein folded structure, the protonation/deprotonation of these side chains are sensitive to pH and can significantly affect the drug binding. Large shifts in p*K*_a_ values are observed for at least one of the active site aspartates of *β*-secretase (p*K*_a_ = 3.5 and 5.2)^4^ and HIV-1 protease (p*K*_a_ = 3.4 - 3.7 and 5.5 - 6.5)^5^ compared to the free aspartic acid in water (p*K*_a_ = 3.8). Experimental and computational studies on HIV-1 protease^6^ and *β*-secretase^7,8^ have demonstrated that pH determines the efficacy of ligand binding to these proteins. Therefore, understanding the pH role on ligand binding to a protein is essential for designing efficiently binding drugs.

Biomolecular modeling^9^ and molecular dynamics^10–12^ (MD) simulations have proven to be indispensable tools to probe and understand protein-drug interactions. Unbiased MD binding simulations^13^ accurately captured the final bound pose of drugs in the binding pocket of the G-protein coupled receptors (GPCR) and identified the dominant binding pathways and the associated barriers. Structure-activity studies and computer-aided drug design are successful in developing commercially available drugs for treating cancers, neurodegenerative diseases (Alzheimer’s,^14^ Parkinson’s^15^), viral diseases (Hepatitis A/C,^16,17^ HIV,^18,19^ avian influenza^16,20,21^) etc.

We used both conventional molecular dynamics (cMD) and constant pH molecular dynamics (cpHMD) simulations to study the unbiased and pH-dependent binding kinetics of guanidinium (Gdm^+^) ions to the hen egg-white lysozyme (HEWL). The active site of HEWL has two acidic residues - Glu35 and Asp52, where Glu35 displays a significant shift in p*K*_a_ value to 6.2 compared to the free Glu in water (p*K*_a_ = 4.3). This shift in p*K*_a_ makes the side chain of Glu35 susceptible to protonation depending on the solution pH and would have a marked effect on Gdm^+^ binding. These features makes HEWL an attractive model system to study the pH effects on ligand binding kinetics.

Guanidinium and guanidine motifs can bind in the active sites of enzymes containing two acidic residues^22,23^ and are of particular interest in drug binding studies^24,25^ as they are playing a critical role in various biological, pharmacological and biochemical processes. While the protein denaturant properties of guanidinium salts in high concentrations (≈ M concentrations) are well documented,^26–31^ submolar and lower concentrations of guanidinium salts are shown to inhibit the activity of proteins such as pancreatic ribonuclease A,^32^ pantetheine hydrolase,^33^ and hen egg-white lysozyme.^22,23^ Besides the enzymatic inhibitory nature of guanidine salts at low concentrations, guanidine and its analogs show chemical and physicochemical features, which are of interest in rational drug design. The resonance stabilized structure of the cation makes Gdm^+^ one of the strongest organic bases with a p*K*_a_ of 13.6. The basicity of the guanidinium moiety can be readily tuned by modifying the accompanying functional groups. ^34^ The tunability and the ability to strongly interact with a wide range of anions such as carboxylates,^35^ phosphates^36^ etc. through hydrogen bond formation and charge pairing, makes guanidine-based molecules excellent candidates for various medical and pharmaceutical applications.

Many currently prescribed pharmaceutically active drugs contain guanidine groups. Antimicrobial drugs such as streptomycin, chlorhexidine, and the antimalarial drug proguanil contain a guanidine moiety. ^37^ TAN-1057D, a bisguanidine isolated from *Flexibacter sp*. acts as an effective antibiotic against methicillin-resistant strains of *Staphylococcus aureus*.^38^ Ptilomycalin A-a tricyclic guanidine isolated from sea sponge shows cytotoxic, anti-fungal, and anti-viral activities.^39^ Guanidine based drugs have also shown considerable promise in inhibiting the activity of proteins implicated in (i) neurodegenerative diseases like Alzheimer’s (*β*-secretase inhibitors), (ii) viral diseases (HIV-1 proteases,^40^ avian flu neuraminidases^16^), (iii) angiogenesis in tumors (acid ceramidase). ^41^ The active site of these target proteins comprises one or more acidic amino acids. Recent studies have shown that guanidine-based drugs specifically bind to acidic residues present at the active site of an enzyme via hydrogen-bonded interactions.^40–42^ Crystal structures of the avian flu neuraminidase inhibitor zanamivir show the inhibitor molecule bridging the active site residues Asp152 and Glu229. ^43^

In this paper, we studied the pH effect on the binding of Gdm^+^ to the HEWL active site using cMD and cpHMD simulations. The advantage of studying Gdm^+^ binding to the HEWL active site is that the binding is fast and occurs on a nanosecond time scale making it feasible to probe the binding mechanism using computer simulations exhaustively. The binding free energy surface (FES) from cMD and cpHMD simulations reveals two dominant Gdm^+^ binding pathways populating multiple intermediate states. We show that pH modulates the population of intermediates in the FES, resulting in strong pH-dependent binding of Gdm^+^ to the active site. The computed association (binding) constant *k*_on_ and the dissociation (unbinding) constant *k*_off_ show a strong pH dependence and a sharp jump in their values when the pH exceeds the p*K*_a_ of Glu35 in the HEWL active site. Analysis of the binding modes reveals the existence of multiple Gdm^+^ bound to the active site indicating the possibility of designing large molecules with multiple guanidinium moieties that can efficiently bind to inhibit enzyme activity. These results provide insight into the pH-dependent binding mechanism of ligands to proteins with acidic residues in the active site and aid in understanding the binding of complex guanidine-based drug molecules to proteins.

## Methods and Materials

### Conventional Molecular Dynamics Simulations

The initial structure for both the conventional (cMD) and constant pH (cpHMD) molecular dynamics simulations is taken from the crystal structure of HEWL (PDB ID: 1AKI).^44^ To study the mechanism of Gdm^+^ binding to the protein, we placed 10 Gdm^+^ randomly around the protein at a minimum cutoff distance of 15 Å from the protein center of mass. The protein and Gdm^+^ are solvated with water molecules in a cubic box of length 80 Å. The concentration of Gdm^+^ in the simulation box is ≈ 32 mM. The excess charge in the simulation box is neutralized by adding 18 Cl^−^ ions. To rule out concentration effects, a second set of simulations with five Gdm^+^ in the simulation box ([Gdm^+^] ≈ 16 mM) were carried out.

We have used the explicit solvent CHARMM36m force field ^45^ for the protein. The parameters for Gdm^+^ are taken from the CGenFF.^46^ Water molecules are modeled using the TIP3P model.^47^ Langevin dynamics simulations are carried out using the NAMD software package (version 2.13)^48^ with periodic boundary conditions applied in all directions. A 12 Å cutoff is used for van der Waals and short-range electrostatic interactions. Long-range electrostatic interactions are calculated using the particle mesh Ewald algorithm.^49^ The bonds involving hydrogen atoms are constrained using the SHAKE algorithm. A 2 fs time step is used to advance the simulation.

Before running the simulations, the system is energy minimized using the conjugate gradient algorithm to remove high-energy contacts. The system is then subjected to a three-step equilibration process: (i) equilibration of water in the *NV T* ensemble for 10 ns with the positions of heavy atoms of the protein and all ions constrained using the fixed atoms algorithm implemented in NAMD. (ii) equilibration of water and chloride ions in the *NPT* ensemble for 10 ns with the positions of heavy atoms of the protein and Gdm^+^ constrained. (iii) In the final step, the positional constraints on heavy atoms of the protein and Gdm^+^ are removed, and the whole system is allowed to equilibrate in the *NPT* ensemble. The temperature *T* is maintained at 300 K using Langevin dynamics, while the pressure *P* is maintained at 1 atm using a Nose-Hoover Langevin piston. We ran 10 independent simulations each of length ≈ 200 ns, and 9 independent simulations each of length ≈ 80 ns in the *NPT* ensemble. For each simulation, the initial configurations of Gdm^+^ are different and are generated by randomly placing the Gdm^+^ in the simulation box. The velocities are randomly generated from the Maxwell-Boltzmann distribution corresponding to *T* = 300 K. We also ran 3 long simulations, each of length 1 *µ*s in addition to the short simulations described above. The extended simulations are carried out using the GROMACS simulation package (version 2020.2).^50,51^ The same forcefield and equilibration protocols described before are used for these simulations as well. In the GROMACS runs, the heavy atoms of the ions and proteins are restrained by applying harmonic restraints using a force constant of 1000 kJ/mol/nm^2^ for the first two equilibration steps as described previously. The temperature is maintained at 300 K using the stochastic velocity rescaling (modified Berendsen) algorithm^52^ implemented in GROMACS with a coupling parameter *τ*_*T*_ = 0.1 ps, and the pressure is maintained at 1 atm using the Parinello-Rahman barostat^53^ with a coupling parameter *τ*_*P*_ = 2.0 ps. The conformations are saved at every 10 ps and the cumulative simulation time is ≈ 5.7 *µ*s. We ran seven independent simulations with lower [Gdm^+^] (≈ 16 mM) using the GROMACS-2020.2 package and CHARMM36m forcefield following the same protocols as described above. The cumulative simulation time for lower [Gdm^+^] is 3.3 *µ*s.

To check that the results are not forcefield dependent, we also ran simulations using the Amber ff99sb-ildn ^54^ forcefield. Gdm^+^ forcefield parameters are obtained using the generalized Amber force field (GAFF)^55^ (Table S1). The charges on atoms in Gdm^+^ are parameterized using the restrained electrostatic potential (RESP) implemented in the antechamber^56^ tool of AmberTools18.^57^ We ran the Amber force field simulations using the GROMACS-2020.2 package following the above-mentioned protocols. We generated three independent trajectories, each of length 600 ns for [Gdm^+^] = 32 mM.

### Constant pH Molecular Dynamics (cpHMD) Simulations

The active site in HEWL consists of two negatively charged residues, Glu35 and Asp52 (Figure 1A). Experimental studies ^58^ show that HEWL is active over the pH range 6 to 9. For HEWL to be in the active state, either one of the two acidic residues should be in a protonated state.^59^ Titration studies on HEWL reveal that the p*K*_a_ values of Glu35 and Asp52 are 6.2 and 3.6, respectively. ^60^ In the active pH range of HEWL, the p*K*_a_ value of Glu35 indicates that it exists in a dynamic equilibrium between the protonated and deprotonated states, whereas Asp52 exists dominantly in its deprotonated form. As Gdm^+^ binds to the active site residues via hydrogen bond formation, the protonation of Glu35 should affect the binding rate of Gdm^+^. To study the effects of Glu35 protonation on Gdm^+^ binding, we setup cpHMD simulations over the pH range 5.0 to 7.0.

**Figure 1:**
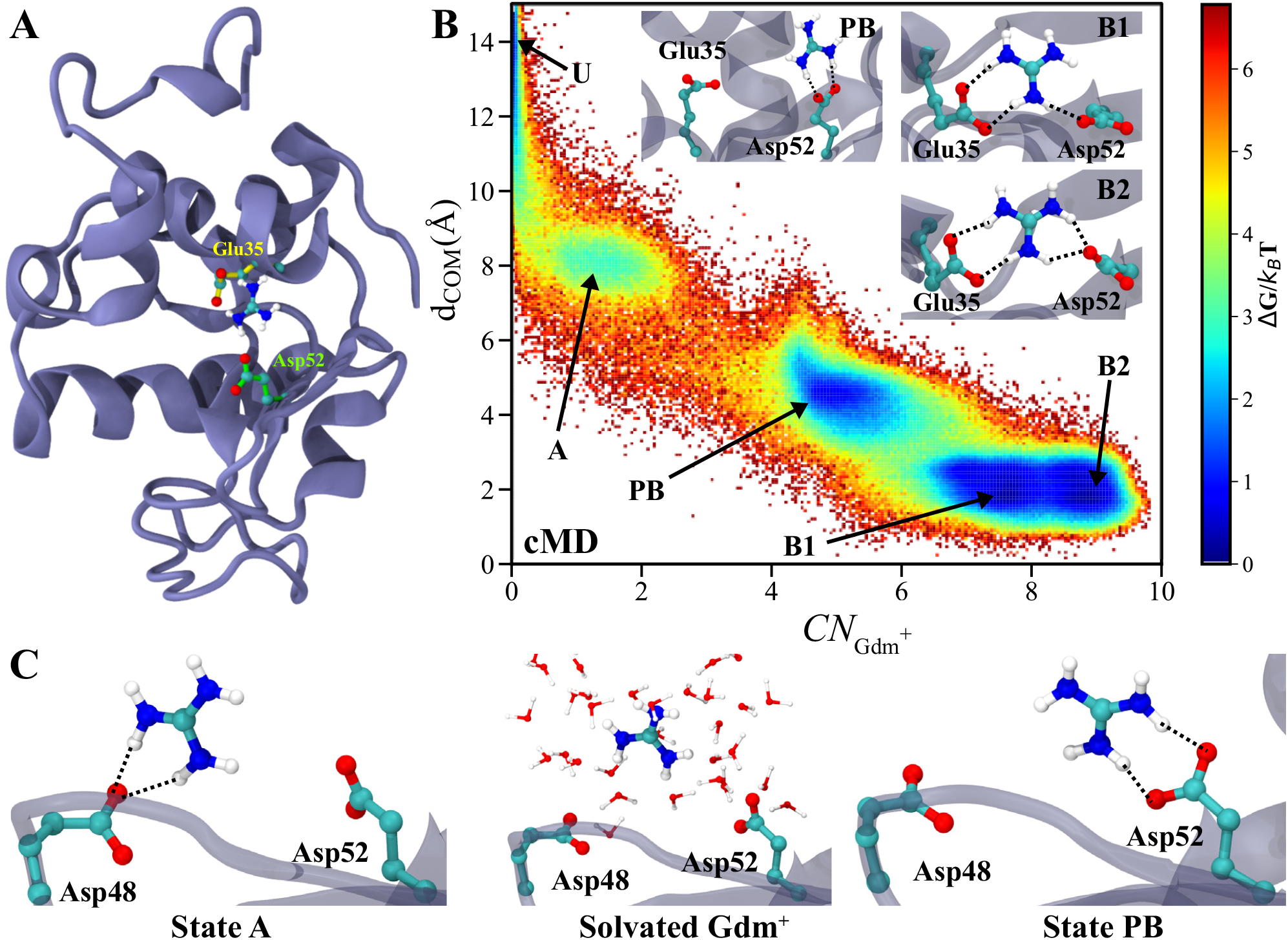
(A) Structure of hen egg-white lysozyme with the active site residues Glu35 (yellow) and Asp52 (green). Gdm^+^ is bound in the active site and interacts with Glu35 and Asp52. The O atoms of the carboxylic groups are shown in red, N atoms of Gdm^+^ are shown in blue, and H atoms of Gdm^+^ are shown in white. (B) The FES of Gdm^+^ binding to the HEWL active site is projected onto 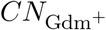 and *d*_COM_. The basins in the FES correspond to the unbound state (**U**), associated state (**A**), partially bound state (**PB**), and the two bound states (**B1** and **B2**). Representative conformations of Gdm^+^ corresponding to the various basins in the FES are shown in the insets. (C) Representative structures show the hopping transition of the Gdm^+^ bound to Asp48 (state **A**) to the active site Asp52 (state **PB**) through the solution.

We performed cpHMD simulations in explicit solvent using the constant pH algorithm implemented in the NAMD software package (version 2.13).^48^ For the explicit solvent cpHMD simulations, we used the CHARMM36m force field^45^ for the protein, CGenFF parameters^46^ for Gdm^+^, and TIP3P water model.^47^ The forcefield also includes the topology and parameters of the protonated states of titratable residues.^45^ In the cpHMD simulations of HEWL, the p*K*_a_ values of the titratable residues present in the protein are obtained from experiments^60^ and are given as an input to the simulations. Cysteine residues involved in the formation of four disulfide bonds in the protein are excluded from the protonation protocol. The simulation protocol we followed for cpHMD is identical to the protocol followed for the cMD simulations. Langevin dynamics simulations are carried out at *T* = 300 K with a time step of 2 fs. Each protonation cycle of the cpHMD simulations comprises 15 ps of non-equilibrium MD switching simulation with 100 ps of conventional MD. We performed cpHMD simulations for [Gdm^+^] = 16 and 32 mM. For [Gdm^+^] ≈ 32 mM, we performed simulations at pH values of 5.0, 5.5, 6.0, 6.5 and 7.0. For each pH value, we generated 3 independent trajectories each of length 500 ns and one longer trajectory of length 1 *µ*s resulting in a cumulative simulation time of 2.5 *µ*s per pH value for [Gdm^+^] = 32 mM. For [Gdm^+^] ≈ 16 mM, we performed simulations at pH values of 6.0, 6.5 and 7.0 with 3 independent trajectories each of length 400 ns resulting in a cumulative simulation time of 1.2 *µ*s per pH value.

### Coordination Number of Gdm^+^ Bound to the Active Site

Gdm^+^ binds to the active site of HEWL by forming a hydrogen-bonded bridge between the carboxylic groups of the negatively charged residues, Glu35 and Asp52 (Figure 1A,B). To monitor the interaction of Gdm^+^ with the active site residues, we calculated the distance (*d*_COM_) between the center of mass of a Gdm^+^ and the center of mass of the active site residues, Glu35 and Asp52. Gdm^+^ is considered bound to the active site, if *d*_COM_ ≤ 3 Å. While *d*_COM_ can indicate that the ion has entered the binding site, it does not provide information about the hydrogen-bonded interactions between Gdm^+^ and carboxylic groups.

To identify the binding pose of a single Gdm^+^, we used the coordination number 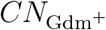 between the carboxylic group O atoms of Glu35 and Asp52, and the N atoms of the Gdm^+^.

The coordination number between the two sets of atoms is calculated using

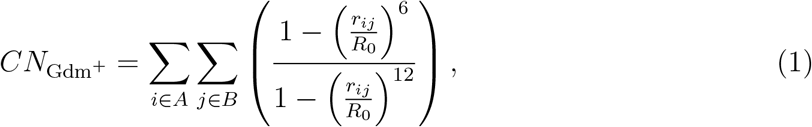

where group A consists of the N atoms of the single Gdm^+^ ion, and group B consists of the carboxylic group O atoms of Glu35 and Asp52, *R*_0_ (= 5 Å) is the cutoff distance and *r*_*ij*_ is the distance between the atoms *i* and *j*. Another CV, 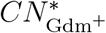, based on the coordination number of all Gdm^+^ in the simulation box with respect to the active site is computed to identify the number of Gdm^+^ binding to the active site. In computing 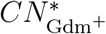, group A in eq. 1 consists of the N atoms of all the Gdm^+^ ions in the simulation box, while group B is the same as in the previous definition. The coordination values are calculated using the COORDINATION function of the PLUMED library. ^61,62^

### Gdm^+^ Binding Free Energy Surface (FES)

We computed the FES associated with the Gdm^+^ binding/unbinding process using Δ*G*(**s**) = -*k*_B_*T* ln(*P* (**s**)), where **s** is the set of collective variable(s) (CVs) describing the process. The CVs used to project the FES are *d*_COM_ and 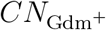. The two-dimensional FES describing the binding process is constructed using the joint probability distribution of *d*_COM_ and 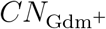, and is given by

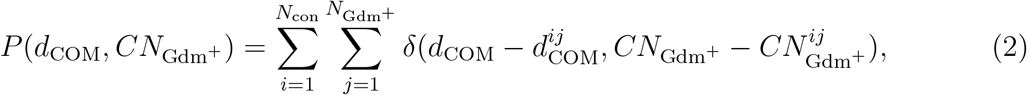

where 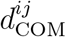 and 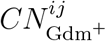 are the *d*_COM_ and 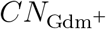, respectively, of the *j*^th^ Gdm^+^ ion in the simulation box in the *i*^th^ simulation snapshot, *N*_con_ is the total number of simulation snapshots, and 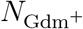 is the total number of Gdm^+^ ions in the box. The one-dimensional FES describing the higher order bound states of Gdm^+^ is constructed using the probability distribution of 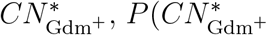.

### Kinetics of Gdm^+^ Binding to the Active Site

A Gdm^+^ is considered bound in the active site if it satisfies: (i) *d*_COM_ < 3 Å and (ii) 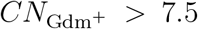. We observed multiple events in the simulations where a bound Gdm^+^ unbinds for a short period and returns to its bound state, or an unbound Gdm^+^ binds for a shorter period and returns to the unbound state. In the analysis, we considered only events where Gdm^+^ remained in the bound/unbound state for at least 0.5 ns as part of a binding/unbinding event.

The Gdm^+^ binding/unbinding rate constants are calculated using the following equations,^63^ which are based on Kramers’ reaction rate theory.^64^ Assuming that Gdm^+^ unbinding follows first-order kinetics, we estimated the unbinding rate constant *k*_off_ using the equation

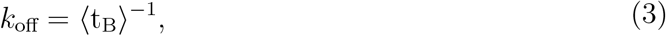

where ⟨t_B_⟩ is the average lifetime of the ligand bound state. Similarly, for the binding process, we assume that the concentration of ligand is in excess over the protein and binding does not significantly alter the ligand concentration. Under these assumptions, the binding rate constant (*k*_on_) is approximated as

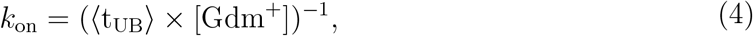

where ⟨t_UB_⟩ is the average lifetime of the unbound state. ⟨t_UB_⟩ and ⟨t_B_⟩ are approximated as the mean first passage time (MFPT) associated with Gdm^+^ binding and unbinding, respectively. In computing ⟨t_UB_⟩ and ⟨t_B_⟩, we ignored events in the simulations where a Gdm^+^ binds or unbinds transiently for short time periods (≤ 0.5 ns).

### Markov State Modeling (MSM)

Simulation data reveals that the HEWL active site can accommodate binding of up to three Gdm^+^ ions in different poses. While the unbound Gdm^+^ in the simulation box undergo diffusion without interacting with the protein’s active site, transient interactions are observed with other acidic residues far away (> 15 Å) from the protein active site. To identify the relevant kinetic processes associated with the binding of multiple Gdm^+^ to the HEWL active site, we constructed a Markov State Model (MSM) using the PyEMMA software package. ^65^ To build the MSM, we used the distances between the following groups of atoms as the input variable: i) heavy atoms of the Gdm^+^ ions, and ii) oxygen atoms of the active site residues - Glu35 (O*ϵ*_1_, O*ϵ*_2_, O) and Asp52 (O*δ*_1_, O*δ*_2_, O), O*δ*_1_ of Asn46 and backbone carbonyl oxygens of Leu56 and Gln57, resulting in a total of 108 distances (Figure S1). MSM construction requires careful consideration of the CVs that capture the dynamics of the relevant process (in this case, binding/unbinding of Gdm^+^). Including the unbound Gdm^+^ in the analysis increased the amount of input data and enhanced the complexity. Hence, we excluded the unbound Gdm^+^ from the analysis and effectively reduced the system to just the protein and three Gdm^+^ ions binding to the active site. To exclude the unbound Gdm^+^ ions from the analysis, we considered only those Gdm^+^, which at any point in the trajectory satisfy *d*_COM_ *<* 5 Å and the rest are ignored.

To identify the relevant microstates in the binding mechanism, we carried out *k*-means clustering^65^ using the computed 108 distances between the relevant atoms of the binding pocket and the Gdm^+^ heavy atoms. To get an initial estimate of the number of clusters required to capture the binding mechanism accurately, we performed *k*-means clustering of the 108 active site-Gdm^+^ distances with 75, 100, 250, 300, 500, 750, 1000, 1250, 1500 and 2000 cluster centers. Given the high dimensionality of the input variable, understanding the significance of these clusters in Gdm^+^-HEWL binding poses a challenge. To get a more intuitive understanding of these clusters, each cluster is projected onto a more relevant 2D space in the context of Gdm^+^ binding. For each cluster obtained from the *k*-means clustering, calculating 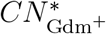 and the average number of hydrogen bonds (⟨N_HB_⟩) between the Gdm^+^ ions and the active site residues Glu35 and Asp52 gave a reasonable understanding of the underlying binding process. The lower number of clusters (<500) showed a poor sampling of the basins associated with the Gdm^+^ binding mechanism while increasing the number of clusters (> 1000) did not significantly alter the distribution of points in the 2D space (Figure S2). We chose 1500 clusters, which is a reasonable number to construct the MSM (Figure S2E). To determine the appropriate lag time to construct the MSM, we calculated the implied timescale as a function of lag time. The implied timescales flattened out at ≈ 1 ns lag time, and we used this value to construct the MSM (Figure S3A). The separation between the first two and the other timescales indicated that MSM resolved the three slow processes corresponding to the free, associated and bound states of the Gdm^+^-HEWL complex. To validate the MSM, we carried out the Chapman-Kolmogorov test (Figure S3B). To understand the kinetics of the multiply bound Gdm^+^ states, the clusters were further coarse-grained based on 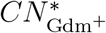 and ⟨N_HB_⟩ (Figure S3C). While the above coarse-graining clearly differentiated between the various bound states, it is unable to discriminate between the unbound state **U** and the associated state **A** (Figure 1B). To separate these two states, we applied an additional criterion, based on the occupancy of a Gdm^+^ near the residue Asp48. If a Gdm^+^ is present within 5 Å of Asp48, we classified the cluster as state **A**, else it is classified as state **U**. This procedure applied to the Gdm^+^-HEWL complex allowed the calculation of the mean first passage times and fluxes between the coarse-grained clusters corresponding to the bound states of Gdm^+^.

## Results and Discussion

### Intermediates States are Populated in Gdm^+^ Binding to the HEWL Active Site

The FES projected onto *d*_COM_ and 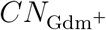 (see Methods) using the data obtained from cMD simulations shows four distinct basins: (i) the unbound state 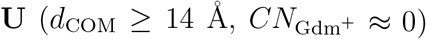, (ii) the associated state **A**, where Gdm^+^ interacts with Asp48, which is present at a distance from the active site residues 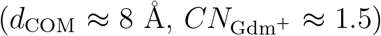, (iii) a partially bound state **PB**, where Gdm^+^ forms hydrogen bonded interactions with only one of the two acidic residues (Asp52 or Glu35) in the active site 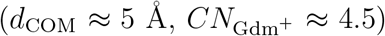, (iv) the bound state **B**, which can be further divided into two substates **B1** 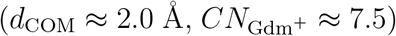 and **B2** 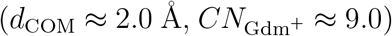 (Figure 1B). The substates **B1** and **B2** are separated by a barrier ≈ 0.9 kcal/mol.

In the first pathway, the state **A** forms an initial entry point for Gdm^+^ to enter the HEWL active site. In this state, a Gdm^+^ interacts with Asp48 using hydrogen-bonded interactions. However, one of the carboxylate oxygens of Asp48 acts as an acceptor in the hydrogen-bond formation with the amide hydrogen and hydroxyl group of neighboring Ser50. As a result, Gdm^+^ can interact with only the other carboxylate oxygen of Asp48 (Figure S4A). The Gdm^+^ also cannot form a bridging interaction between Asp48 and active site Asp52 as the average distance between the carboxylate groups of these two residues (≈ 9.0 Å) is larger than the ideal distance (≈ 7 Å) required for a Gdm^+^ to anchor itself between these residues.^23^ However, compared to Asp48, the nearby active site residues Glu35 and Asp52 have their carboxylate oxygens in an orientation favorable to form hydrogen-bonded interactions with Gdm^+^ resulting in a more favorable interaction. As a result, Gdm^+^ hops through the solvent from Asp48 to the active site Asp52 populating either the **PB** or **B1/B2** state in the FES. The state **B1/B2** is more stable due to the additional hydrogen-bonded interactions over those present in state **PB** (Figure 1C and S4B). In this pathway, when the **PB** state is populated, Gdm^+^ interacts with only the Asp52 carboxylate group in the active site (Figure 1C). The other possibility, i.e., Gdm^+^ interacting with Glu35 in the active site, is not observed.

In a different Gdm^+^ binding pathway, the state **A** is not populated. In this pathway, Gdm^+^ from the solution directly interacts with Glu35 in the active site populating the **PB** state (Figure S5). The probability of observing this pathway is low because compared to Asp52, which is solvent exposed, Glu35 is partly occluded by residues Trp108 and Arg114, blocking its direct access to Gdm^+^ from the solvent. The residue Arg114, in particular, plays a vital role in preventing Gdm^+^ from binding to Glu35 in state **PB** (Figure S6A,B). As the amino acid arginine has a guanidinium moiety in its side chain, Arg114 can occupy a position similar to that of a bare Gdm^+^ in the protein’s active site. However, due to conformational constraints, Arg114 can only form hydrogen bonds with Glu35. The distance between Glu35 and Arg114 in state **PB** shows that Arg114 is hydrogen-bonded to Glu35, preventing direct access of Glu35 to Gdm^+^ (Figure S6A,B).

In the bound state **B**, a single Gdm^+^ interacts through hydrogen bonds between its N atoms and carboxylate groups of Glu35 and Asp52. As each of the three N atoms of Gdm^+^ can independently act as hydrogen bond donors to the active site carboxylate oxygens, relative orientations of both active site residues and the Gdm^+^ can lead to several combinations of hydrogen-bonded interactions (Figure S7). Thermal fluctuations in the active site can change donor-acceptor distances while keeping the hydrogen bond interactions intact. The collective variable 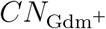 is sensitive to the distances between the selected groups (Eq. 1). Even small changes in the donor-acceptor distances due to thermal fluctuations between the bound Gdm^+^ and active site carboxylate groups can lead to a significant difference in the CN value (Figure S8), which leads to a broad **B** state basin along the CN axis in the FES (Figure 1B). The multiple bound poses of Gdm^+^ in the broadly distributed **B** state contribute to the substates, **B1** and **B2** in the FES (Figure 1B and S7). The substate **B2** is more stable than **B1** due to the formation of stronger hydrogen bonds as the donor-acceptor distances are optimum, which is also supported by the increase in 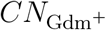.

### pH Effect on the Population of Intermediate States

The protonation state of the carboxylic groups of Glu35 and Asp52 depends on the residue p*K*_a_ values and the solution pH. The protonation state of the residues in the active site strongly influences the Gdm^+^ binding to the active site. The p*K*_a_ values of Glu35 and Asp52 are 6.2 and 3.6, respectively, ^60^ and HEWL is active over a pH range (pH = 6 - 9).^58^ In the active pH range, Asp52 is predominantly deprotonated, while Glu35 can have a variable protonation state dependent on the solution pH. The percentage protonation of Glu35 and Asp52 in the active pH range is calculated using the Henderson-Hasselbach equation. At pH = 5.0, ≈ 94% of Glu35 and ≈ 3.8% of Asp52 is protonated in bulk, whereas at pH = 7.0, the bulk protonation of Glu35 and Asp52 are ≈ 13.7% and ≈ 0.04%, respectively. A titratable residue is considered to be roughly 99.9% deprotonated at a pH value 2 units above the p*K*_a_. In the cMD simulations, as both Glu35 and Asp52 are deprotonated, we approximate the pH value of these cMD simulations is ≈ 8.5. Gdm^+^ binding to the HEWL active site depends on the hydrogen bond formation between the N atoms of Gdm^+^ and the carboxylic group O atoms of Glu35. The protonation of Glu35 makes it unable to act as a hydrogen bond acceptor, thereby preventing the formation of hydrogen bonds and strong binding of Gdm^+^ in the active site. However, deprotonation of both acidic active site residues at high pH (≈ 8.5) (Figure 1B) results in strong binding of Gdm^+^ (Figure 1B).

At pH = 5.0, ≈ 94% of Glu35 is protonated and cannot interact via hydrogen bond formation with Gdm^+^. However, the Asp52 side chain is predominantly deprotonated and can form hydrogen bonds with Gdm^+^, making the **PB** state the most stable Gdm^+^-active site interaction in the FES (Figure 2A). As the pH increases, the concentration of the deprotonated state of Glu35 increases, and Gdm^+^ can form the hydrogen-bonded bridge between Glu35 and Asp52, which results in the appearance of the singly-bound state **B** at pH = 6.0 (Figure 2B). At pH = 6.5, the percentage of deprotonated Glu35 is ≈ 66.6%, which allows Gdm^+^ to properly orient itself between the active site residues and sample multiple bound poses with different combinations of hydrogen bonds resulting in the appearance of the broad **B** state basin (Figure 2C). At a higher pH value of 7.0, the basin corresponding to the bound state becomes stable and splits into two substates, **B1** and **B2**, where *d*_COM_ is ≈ 2 Å for both the states but 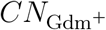 value is ≈ 7.5 and ≈ 9.0, respectively (Figure 2D).

**Figure 2:**
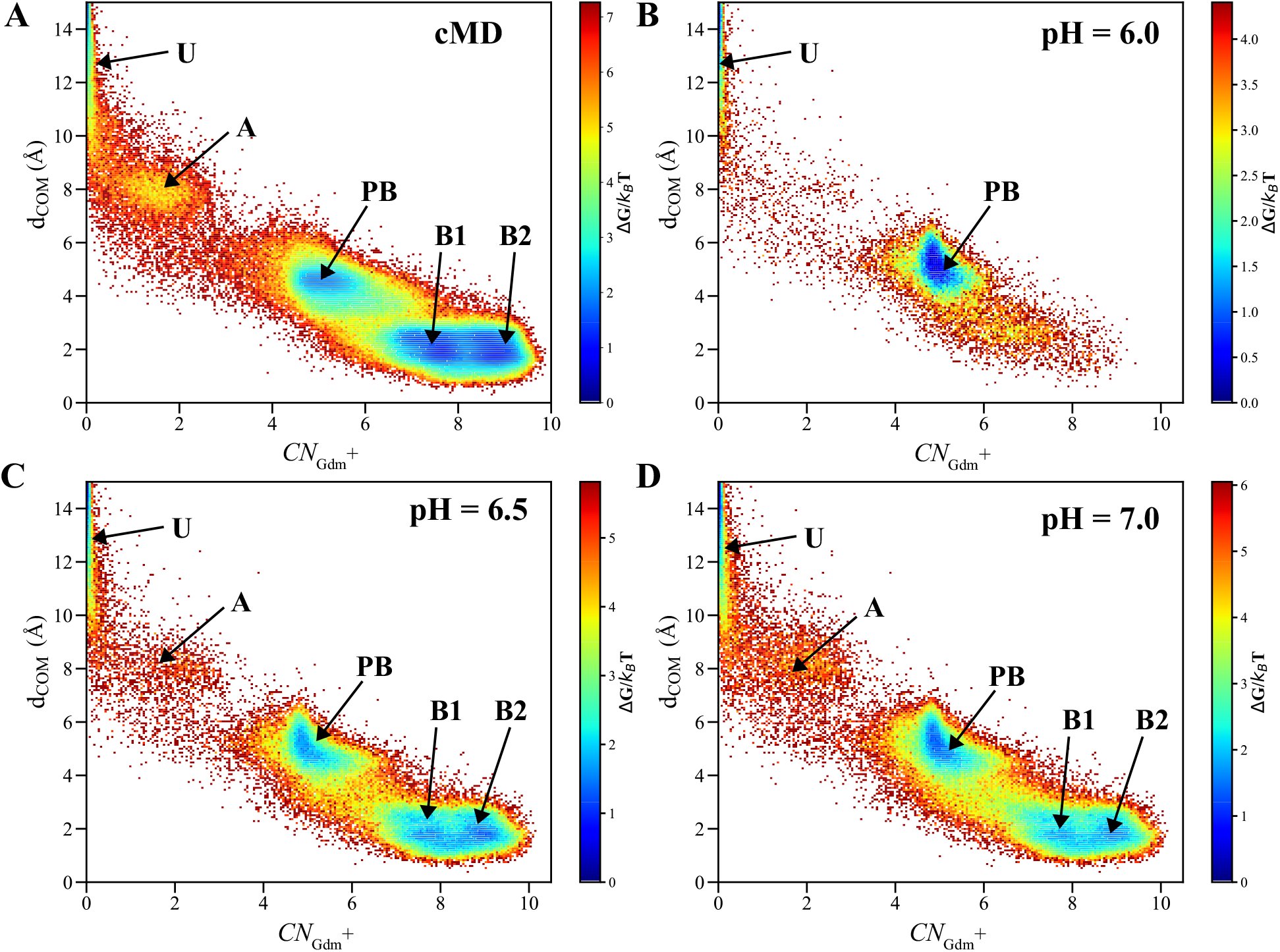
FES projected onto 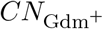 and *d*_COM_ for different pH values. The minima marked on the FES correspond to different Gdm^+^ binding states. At low pH, the high protonation rate of Glu35 contributes to the absence of the bound state basins, **B1** and **B2**. At higher pH, the Gdm^+^ bound state is the global minimum.

### Multiple Gdm^+^ Can Bind to the HEWL Active Site

The simulation data shows that while a Gdm^+^ is forming bridging interactions between the carboxylic residues in the HEWL active site, it can act as a scaffold for further formation of hydrogen-bonded interactions between the active site residues and another Gdm^+^.^23^ To identify these complexes, we computed the coordination number 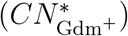 of the N atoms of all the Gdm^+^ present in the simulation box with the carboxylic group O atoms of the active site residues Glu35 and Asp52 (see Methods) (Figure 3).

**Figure 3:**
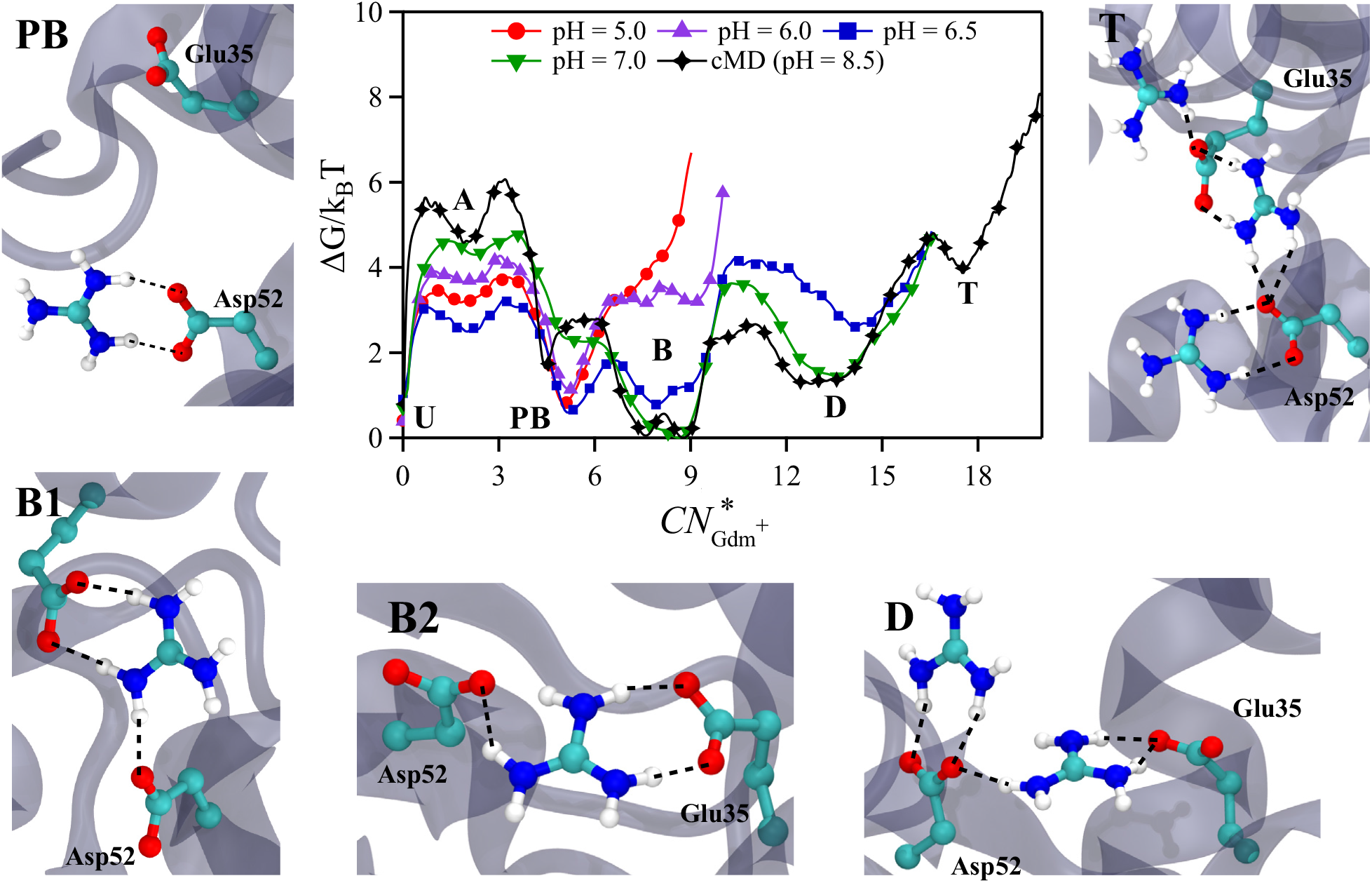
FES projected onto 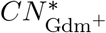. The minima corresponding to the various configurations of Gdm^+^ are marked on the FES. States **D** and **T** denote the double and triply bound states of Gdm^+^ to the active site, respectively. Increasing the pH leads to an increased stability of the singly bound state **B**, which in turn enhances the stability of state **D**. The insets depict representative structures of the bound states of Gdm^+^.

The FES projected onto 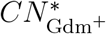 reveals multiple minima corresponding to the different states in Gdm^+^ binding to the HEWL active site (Figure 3). The minima in the FES correspond to: (i) unbound state 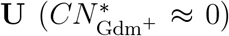, (ii) the associated state **A** where the Gdm^+^ is bound to Asp48 in the vicinity of the active site 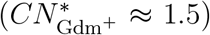, (iii) the partly bound state **PB**, where a Gdm^+^ is hydrogen-bonded to a single carboxylic group in the active site (either Glu35 or Asp52) 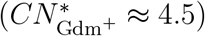, (iv) the singly-bound state **B**, which is further separated into two shallow basins **B1** and **B2**, (v) the bridged state **D**, where one Gdm^+^ forms the hydrogen-bonded bridge between the carboxylic groups of Glu35 and Asp52 and another Gdm^+^ is hydrogen-bonded to the Asp52 carboxylic group 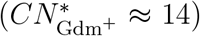 and (vi) the doubly bridged state **T**, where one Gdm^+^ forms the hydrogen-bonded bridge between the carboxylic groups of Glu35 and Asp52 and two other Gdm^+^ ions are hydrogen-bonded to the Glu35 and Asp52 carboxylic groups, respectively 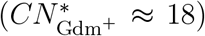 (Figure 3). The FES projected onto 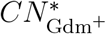 for different pH values at lower [Gdm^+^] exhibit similar minima (with the exception of state **T**), indicating multiple Gdm^+^ bound to the active site of HEWL is not an artifact of the Gdm^+^ concentration used (Figure S11).

The formation of these higher-order Gdm^+^ complexes is facilitated by the initial bridging interaction between a Gdm^+^ and the active site residues. As the bridge formation depends upon the pH of the system, we computed the FES projected onto 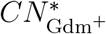 for different pH (Figure 3). As the binding of the second Gdm^+^ would depend on the local concentration of Gdm^+^ around the protein, we calculated the radial distribution function (*g*(*r*)) of the center of mass of Gdm^+^ around the carboxylate groups of active site residues Glu35 and Asp52 for different pH values (Figure S12). At pH = 5, the state **PB** is predominantly populated in the FES, while the other states-**B1, B2, D** and **T** are almost absent. This is also evident from the *g*(*r*) of Gdm^+^ around the active site residues (Figure S12). The height of the primary peak in *g*(*r*) at ≈ 3 Å decreased with the decrease in pH, indicating that the local concentration of Gdm^+^ in the vicinity of the active site residues decreased. The peak corresponding to the presence of a secondary Gdm^+^ (≈ 5 Å) is absent at low pH, indicating the absence of higher-order bound states, **D** and **T**. This implies that at pH smaller than the p*K*_a_ of Glu35, neither the singly-bound nor higher-order bound states are stable, and protonation of Glu35 is more favorable than forming hydrogen bond bridges with Gdm^+^.

Upon increasing the pH to 6.5, the minima depth corresponding to the **B** state increased, indicating that the hydrogen-bonded network between a Gdm^+^ and Glu35 is stable due to the deprotonation of Glu35. The height of the primary peak in *g*(*r*) at ≈ 3 Å also increased, mediated by a stronger interaction between Gdm^+^ and the active site residues. The presence of another peak at ≈ 5 Å in the *g*(*r*) indicates the binding of a secondary Gdm^+^ in the vicinity of the binding site leading to the formation of an additional non-bridging hydrogen bonded interaction (Figure S12). Since the stability of the bridging interaction in the single bound state of Gdm^+^ dictates the formation of higher-order Gdm^+^ complexes, we see the population of the doubly bound state even at pH = 6.5, which is slightly above the p*K*_a_ value of Glu35 (p*K*_a_= 6.2) (Figure 3).

### Gdm^+^ Binding Rate Constants Depend on the pH

We computed the binding (*k*_on_) and unbinding (*k*_off_) rate constants (Eq. 3 and 4) of Gdm^+^ to the HEWL active site using the data from both the cMD and the cpHMD simulations (Figure 4A). At pH equal to the p*K*_a_ of Glu35 (p*K*_a_= 6.2), there is a jump in the *k*_on_ values. At lower pH, only Asp52 in the active site can interact with a Gdm^+^, which decreases the overall stability of the binding complex, and Gdm^+^ also unbinds relatively faster from the active site as the *k*_off_ values are higher (Figure 4A). At pH greater than the p*K*_a_ of Glu35, increased deprotonation of the Glu35 leads to both acidic residues forming hydrogen-bonded interactions with a Gdm^+^, resulting in higher *k*_on_ values and also lower *k*_off_ values. The same effect is observed in the cMD simulations (pH ≈ 8.5), where both the acidic residues are deprotonated over the entire duration of the simulation, and Gdm^+^ binding is highly favored compared to the unbound state.

**Figure 4:**
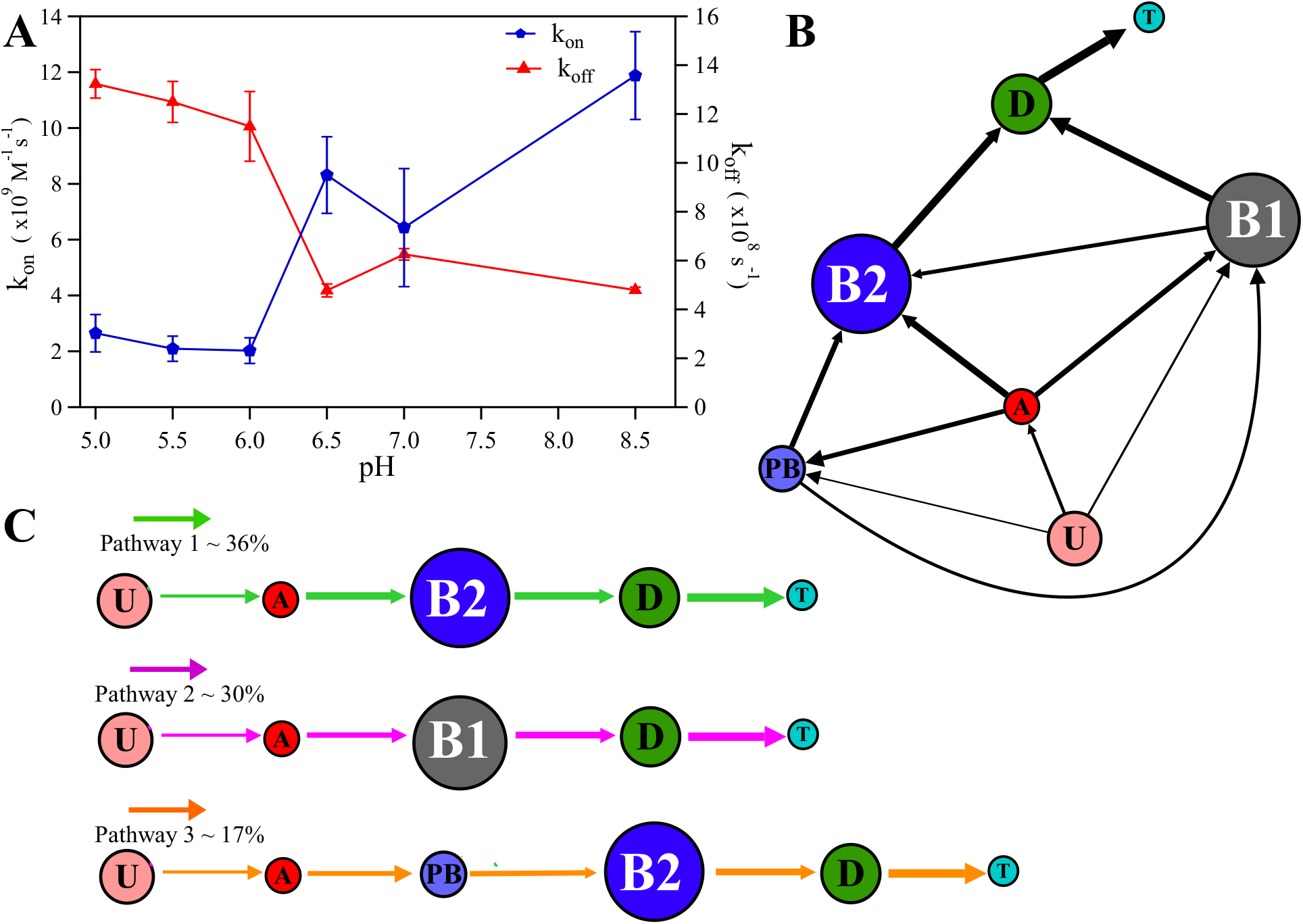
(A) The binding/unbinding rate constants plotted for different pH values. (B) The network of probability fluxes depicting transitions between state **U** and **T** of the Gdm^+^-protein complexes obtained from the MSM. The MSM has been constructed using the data from the cMD simulations (pH ≈ 8.5). The size of the circles indicates the equilibrium population of each state, while the black arrows indicate the fluxes between the states. The thickness of the arrows indicates the amount of flux passing between the states. (C) The fluxes passing through the dominant pathways Pathway 1, Pathway 2 and Pathway 3 are shown with green, magenta and orange arrows respectively.

### Network of Transitions in Gdm^+^ Binding

The existence of higher-order bound states of Gdm^+^, with multiple Gdm^+^ ions occupying the binding pocket, raises questions about the relative stability and the associated kinetic network connecting these states. To elucidate the complex nature of the protein-Gdm^+^ binding/unbinding process, we constructed a Markov State Model (MSM) to extract the relevant kinetic information from the cMD simulations. We analyzed the resulting MSM with the transition-path theory^66,67^ formalism and extracted the transition pathways and the associated timescales (mean first passage time (MFPT)) between the various protein-Gdm^+^ complexes.

From the MSM, we obtained the kinetic network linking the unbound state **U** with the various bound Gdm^+^-protein complexes. From the equilibrium probability distribution, we see the states **B1** and **B2** being the most populated (equilibrium probability *π*_**B1**_ = 0.33 and *π*_**B2**_ = 0.36) and state **T** being the least populated (*π*_**T**_ = 0.007). Direct Gdm^+^ binding, i.e. the transition from state **U** to either state **B1** or **B2**, has a low forward (binding) MFPT_on_ of ≈ 4.4 ns, with the reverse (unbinding) MFPT_off_ (states **B1/B2** to state **U**) of ≈ 43 ns. The associated rate constants can be extracted using Eqns. 3 and 4. The rate constants computed from the MFPT (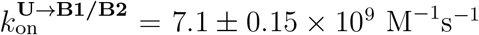 and 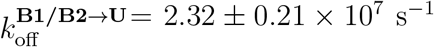) indicate that Gdm^+^ binding/unbinding to the active site are fast processes.

However, the existence of the associated **A** and partly bound **PB** states shows that Gdm^+^ binding from solution to the active site is through the population of intermediates. The conversion timescale from state **PB** to either state **B1/B2** is a fast process (MFPT ≈ 3.5 ns), as it is more energetically favorable for the Gdm^+^ to form the bridged hydrogen-bonded network with both acidic residues rather than just with Asp52. Conversely, the high MFPT from state **B1/B2** to state **PB** (≈ 69 and 84 ns, respectively) shows that Gdm^+^ prefers to be in the bound state interacting with both Glu35 and Asp52. On the transition from state **B1/B2** to **PB** there is a loss of energetic stabilization offered by the formation of two hydrogen bonds on average to the Gdm^+^-active site complex in the state **B1/B2**.

The higher-order complexes, **D** and **T**, show a marked difference in populations with state **D** having a relatively high population (equilibrium probability *π*_**D**_ = 0.23) compared to state **T**. These higher-order states can be considered as composite states, comprised of a mixture of lower-order states (**PB, B1, B2**) as seen from the network of transitions. From the configuration of Gdm^+^ ions in state **D**, it is a composite state comprising of state **PB** and **B1/B2**, with one Gdm^+^ bridging Glu35 and Asp52, and the other tethered to Asp52. State **T** comprises of three Gdm^+^ ions, with two adopting the pose as seen in state **D**, while the third Gdm^+^ is hydrogen-bonded to Glu35. In contrast to the interaction observed between Arg114 and Glu35 in state **PB**, the distance between Arg114 and Glu35 shows that Arg114 is flipped away from Glu35 (≈ 10 Å) (Figure S6C,D), making it feasible for a third Gdm^+^ ion to come and bind to Glu35.

The kinetic pathways show that the state **A** receives the majority of the flux from state **U**, which acts as the gateway for all subsequent bound states (Figure 4B). States **PB** and **B1** also receive a small amount of the flux from state **U**, highlighting the minor pathways comprising of direct Gdm^+^ interactions with Glu35, bypassing the state **A**. From the kinetic network, three dominant pathways are observed linking state **T** to state **A** (Figure 4C). The most dominant pathway, linking states **A, B2, D** to state **T**, comprises roughly 36% of possible pathways. A second less dominant pathway, linking state **A** to **T** via states **B1** and **D** comprises another roughly 30% of possible pathways. A third pathway, making up 17% of the paths, links **A** to **T** via the states **PB, B2** and **D**. The three major pathways described above account for roughly 83% of the total probability flux from state **U** to **T**, with the remaining flux distributed over pathways made up of combinations of these states. In all the observed pathways, state **D**, consisting of two Gdm^+^ in the active site, forms the final stage before forming state **T**. The kinetic network further highlights the importance of the bound state **(B1/B2)** in stabilizing the higher order bound complexes. The higher 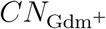 in state **B2** compared to state **B1** is due to an increased stability of hydrogen bonded interactions in **B2**. The higher flux through the state **B2** indicates the relative stability of **B2** over **B1**, and the state **B2** contributes more to the formation and stabilization of states **D** and **T**.

### Gdm^+^ Unbinding Pathways from the HEWL Active Site

As the pharmacological activity of a drug persists until the drug remains bound to the target protein, understanding the drug dissociation pathways and the associated lifetimes or the residence times of the drug becomes a key determinant of pharmacological activity.^68–70^

We show that Gdm^+^ unbinds from the HEWL active site using two distinct pathways. These pathways are dependent on the number of Gdm^+^ bound to the active site. As discussed, Gdm^+^ binding to the active site facilitates further binding of an additional one to two Gdm^+^ to the active site. The presence of these secondary Gdm^+^ in the active site plays an important role in the primary unbinding pathway (Figure 5). This pathway proceeds through the following steps: (i) a 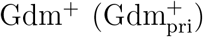) binds to the active site in states **B1** or **B2**, (ii) the binding of 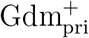 further facilitates the binding of a second 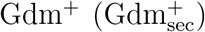) to the active site. The 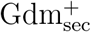 is in a state **PB** and interacts with Asp52. The active site-Gdm^+^ interactions are mainly hydrogen bonds, and thermal fluctuations can disrupt these interactions, leading to a weakening of the hydrogen-bonded network between the active site carboxylates and 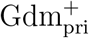. The weakening of the 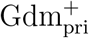 interactions leads to 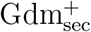 moving into the active site, as seen from the corresponding increase in 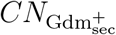 (Figure 5). As a result, the 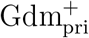 is no longer present in the active site in state **B1/B2**, but moves to state **PB**, being stabilized by the hydrogen bonds with either residue Asp52 or Glu35. However, Gdm^+^ interaction with Glu35 is not stable due to steric clashes as Glu35 is partly occluded by residues Trp108 and Arg114, resulting in Gdm^+^ moving into the solution. A similar scenario is observed in the case of 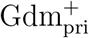 interacting with Asp52. As only one carboxylate moiety is available for the Gdm^+^ to interact with, these interactions are unstable and can be disrupted by thermal fluctuations, driving the Gdm^+^ into solution.

**Figure 5:**
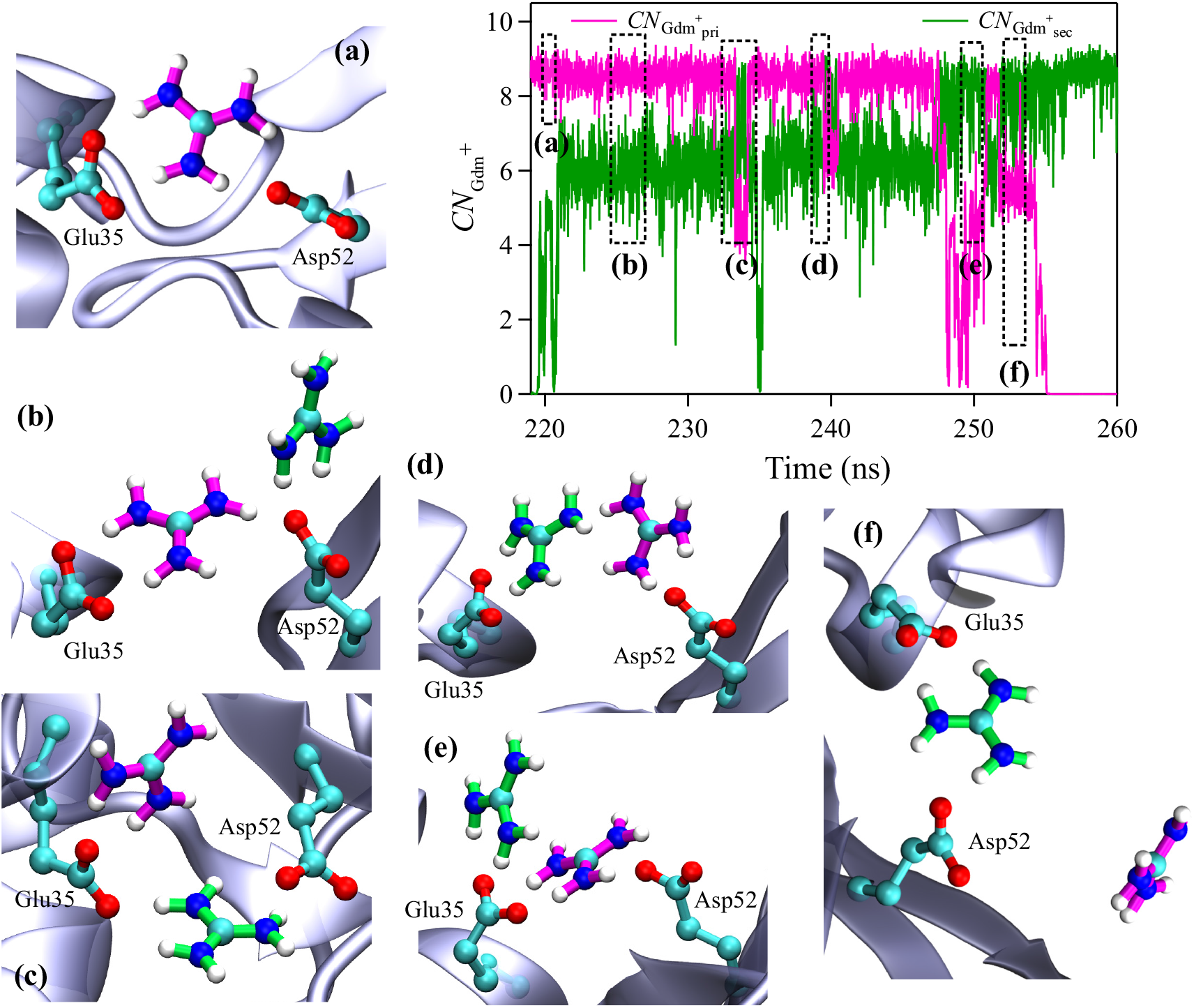
Primary Gdm^+^ dissociation pathway from the active site of HEWL. The coordination number of 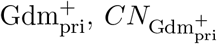 (magenta) decreases as 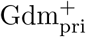 moves away from the state **B1/B2** while the 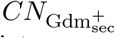 (green) of 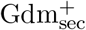 increases as it moves from the state **PB** to **B1/B2**. The insets depict representative structures of key transitions in the unbinding pathway.

The second minor unbinding pathway involves the unbinding of a single Gdm^+^ bound to the active site of HEWL (Figure S13). In this pathway, a Gdm^+^ binds to the active site in state **B1** or **B2**. However, Glu35 is hydrogen-bonded to the guanidine group of Arg114, which destabilizes the active site geometry by twisting the carboxylate group of Glu35 away from the active site, resulting in the disruption of Gdm^+^-Glu35 hydrogen bonds. The breaking of Gdm^+^-Glu35 hydrogen bonds causes the Gdm^+^ to go from state **B1/B2** to state **PB**, with the Gdm^+^ being stabilized by Glu35. As the Gdm^+^ is bound to only one residue, thermal fluctuations can break the hydrogen-bonded interactions, driving the Gdm^+^ into solution.

## Conclusions

Drugs bearing guanidine moieties are becoming increasingly common, from antimicrobial compounds to anti-cancer drugs. However, the binding affinity of these drugs can be sensitive to pH. In this study, we probed the pH dependence of protein-ligand interactions using hen egg-white lysozyme and guanidinium (Gdm^+^) ions as a model system. Gdm^+^ binds to HEWL via hydrogen-bonded bridges to the active site residues, Glu35 and Asp52. The binding free energy surface revealed several basins corresponding to the configurations of Gdm^+^ ions in the active site. At lower pH, Gdm^+^ binding to the active site is destabilized due to increased protonation of Glu35, thereby disrupting the formation of hydrogen bonds between the Gdm^+^ ion and Glu35. Higher pH values stabilize the singly-bound state of Gdm^+^ as well as the formation of secondary non-bridging hydrogen bonds between Gdm^+^ and Asp52.

The computed Gdm^+^ binding and unbinding rate constants show that at low pH values, binding rates decrease while unbinding rates increase. The increased unbinding rate is due to the reduced stability of the Gdm^+^-active site complex, as only one acidic residue (Asp52) can participate in the hydrogen-bonded network. Higher pH values show increased binding rates due to the deprotonation of Glu35, which can then participate in the hydrogen-bonded network, thereby increasing the stability of the Gdm^+^-active site complex.

Previously studies ^22^ showed that multiple Gdm^+^ ions could bind to the HEWL active site. The single-bound Gdm^+^ bridges the two active site residues and acts as a scaffold for one or two additional Gdm^+^ ions to form non-bridging hydrogen bonds with the active site residues. The free energy surface revealed the existence of multiple minima, which corresponds to the states with two and three Gdm^+^ ions bound to the HEWL active site. Markov State Modelling showed the network of transitions between the various bound states and the associated equilibrium probabilities, which showed that the singly-bound state was the most stable state, with the two bound Gdm^+^ states also being appreciably populated.

## TOC Figure

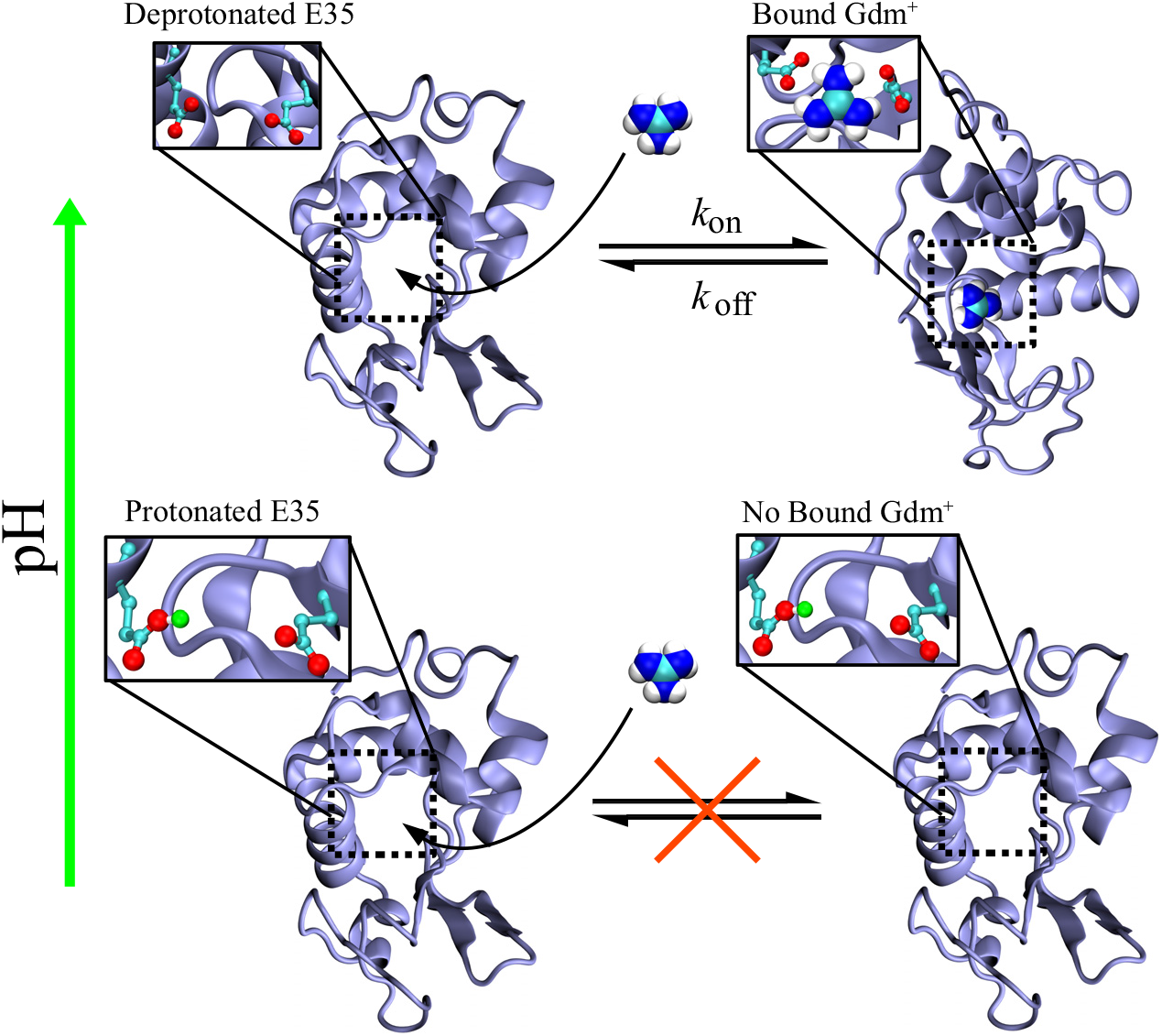

## Notes

### Competing Interest Statement

The authors have declared no competing interest.

## References

(1) Shimizu, H.; Tosaki, A.; Kaneko, K.; Hisano, T.; Sakurai, T.; Nukina, N. Crystal structure of an active form of BACE1, an enzyme responsible for amyloid beta protein production. Mol. Cell. Biol. 2008, 28, 3663–3671.

(2) Ido, E.; Han, H.; Kezdy, F.; Tang, J. Kinetic studies of Human-Immunodeficiency-Virus Type-1 protease and its active-site hydrogen-bond mutant A28S. J. Biol. Chem. 1991, 266, 24359–24366.

(3) Mountford, C.; Grossman, G.; Holmes, K.; O’Sullivan, W.; Hampson, A.; Raison, R.; Webster, R. Effect of monoclonal anti-neutamiidase antibodies on the kinetoc-behaviour of influenza-virus neuraminidase. Mol. Immunol. 1982, 19, 811–816.

(4) Hyland, L.; Tomaszek, T.; Meek, T. Human immunodeficiency virus-1 protease: use of pH rate studies and solvent kinetic isotope effects to elucidate details of chemical mechanism. Biochemistry 1991, 30, 8454–8463.

(5) Toulokhonova, L.; Metzler, W.; Witmer, M.; Copeland, R.; Marcinkeviciene, J. Kinetic studies on β-Site amyloid precursor protein-cleaving enzyme (BACE) - Confirmation of an iso-mechanism. J. Biol. Chem. 2003, 278, 4582–4589.

(6) Gossas, T.; Danielson, U. Analysis of the pH-dependencies of the association and dissociation kinetics of HIV-1 protease inhibitors. J. Mol. Recognit. 2003, 16, 203–212.

(7) Kim, M. O.; Blachly, P. G.; McCammon, J. A. Conformational dynamics and binding free energies of inhibitors of BACE-1: From the perspective of protonation equilibria. PLoS Comput. Biol. 2015, 11.

(8) Kocak, A.; Erol, I.; Yildiz, M.; Can, H. Computational insights into the protonation states of catalytic dyad in BACE1-acyl guanidine based inhibitor complex. J. Mol. Graph. 2016, 70, 226–235.

(9) Jorgensen, W. L. Foundations of biomolecular modeling. Cell 2013, 155, 1199–1202.

(10) Durrant, J. D.; McCammon, J. A. Molecular dynamics simulations and drug discovery. BMC Biol. 2011, 9.

(11) Borhani, D. W.; Shaw, D. E. The future of molecular dynamics simulations in drug discovery. J. Comput.-Aided Mol. Des. 2012, 26, 15–26.

(12) Liu, X.; Shi, D.; Zhou, S.; Liu, H.; Liu, H.; Yao, X. Molecular dynamics simulations and novel drug discovery. Expert. Opin. Drug Discov. 2018, 13, 23–37.

(13) Dror, R. O.; Pan, A. C.; Arlow, D. H.; Borhani, D. W.; Maragakis, P.; Shan, Y.; Xu, H.; Shaw, D. E. Pathway and mechanism of drug binding to G-protein-coupled receptors. Proc. Natl. Acad. Sci. U. S. A. 2011, 108, 13118–13123.

(14) Kumar, S.; Chowdhury, S.; Kumar, S. In silico repurposing of antipsychotic drugs for Alzheimer’s disease. BMC Neurosci. 2017, 18.

(15) Vancraenenbroeck, R.; De Raeymaecker, J.; Lobbestael, E.; Gao, F.; De Maeyer, M.; Voet, A.; Baekelandt, V.; Taymans, J.-M. In silico, in vitro and cellular analysis with a kinome-wide inhibitor panel correlates cellular LRRK2 dephosphorylation to inhibitor activity on LRRK2. Front. Molec. Neurosci. 2014, 7.

(16) von Itzstein, M. et al. Rational design of potent sialidase-based inhibitors of influenzavirus replication. Nature 1993, 363, 418–423.

(17) Njoroge, F. G.; Chen, K. X.; Shih, N.-Y.; Piwinski, J. J. Challenges in modern drug discovery: A case study of boceprevir, an HCV protease inhibitor for the treatment of hepatitis C virus infection. Acc. Chem. Res. 2008, 41, 50–59.

(18) Hdoufane, I.; Bjij, I.; Soliman, M.; Tadjer, A.; Villemin, D.; Bogdanov, J.; Cherqaoui, D. In Silico SAR Studies of HIV-1 Inhibitors. Pharmaceuticals 2018, 11.

(19) Wlodawer, A.; Vondrasek, J. Inhibitors of HIV-1 protease: A major success of structure-assisted drug design. Annu. Rev. Biophys. Biomolec. Struct. 1998, 27, 249–284.

(20) Lew, W.; Chen, X.; Kim, C. Discovery and development of GS 4104 (oseltamivir): An orally active influenza neuraminidase inhibitor. Curr. Med. Chem. 2000, 7, 663–672.

(21) Clark, D. E. What has computer-aided molecular design ever done for drug discovery? Expert Opin. Drug Discov. 2006, 1, 103–110.

(22) Biswas, B.; Muttathukattil, A. N.; Reddy, G.; Singh, P. C. Contrasting effects of guanidinium chloride and urea on the activity and unfolding of lysozyme. ACS Omega 2018, 3, 14119–14126.

(23) Muttathukattil, A. N.; Srinivasan, S.; Halder, A.; Reddy, G. Role of guanidinium-carboxylate ion interaction in enzyme inhibition with implications for drug design. J. Phys. Chem. B 2019, 123, 9302–9311.

(24) Saczewski, F.; Balewski, L. Biological activities of guanidine compounds. Expert Opin. Ther. Patents 2009, 19, 1417–1448.

(25) Zarate, S. G.; Santana, A. G.; Bastida, A.; Revuelta, J. Synthetic approaches to heterocyclic guanidines with biological activity: An update. Curr. Org. Chem. 2014, 18, 2711–2749.

(26) England, J. L.; Haran, G. Role of solvation effects in protein denaturation: from thermodynamics to single molecules and back. Annu. Rev. Phys. Chem. 2011, 62, 257–277.

(27) Tanford, C.; Kawahara, K.; Lapanje, S. Proteins in 6M guanidine hydrochloride - demonstration of random coil behavior. J. Biol. Chem. 1966, 241, 1921–3.

(28) Makhatadze, G.; Privalov, P. Protein interactions with urea and guanidinium chloride - a calorimetric study. J. Mol. Biol. 1992, 226, 491–505.

(29) Lim, W. K.; Rosgen, J.; Englander, S. W. Urea, but not guanidinium, destabilizes proteins by forming hydrogen bonds to the peptide group. Proc. Natl. Acad. Sci. U. S. A. 2009, 106, 2595–2600.

(30) Greene, R.; Pace, C. Urea and guanidine-hydrochloride denaturation of ribonuclease, lysozyme, α-chymotrypsin, and β-lactoglobulin. J. Biol. Chem. 1974, 249, 5388–5393.

(31) Canchi, D. R.; Garcia, A. E. Cosolvent Effects on Protein Stability. Annu. Rev. Phys. Chem. 2013, 64, 273–293.

(32) Miller, J. F.; Bolen, D. A guanidine hydrochloride induced change in ribonuclease without gross unfolding. Biochem. Biophys. Res. Commun. 1978, 81, 610–615.

(33) Pitari, G.; D’Archivio, A. A.; Leandro, L. D.; Antonini, G.; Panatta, A.; Tettamanti, E.; Duprè, S.; Malatesta, F. Conformational changes at the active site of pantetheine hydrolase during denaturation by guanidine hydrochloride. J. Protein Chem. 1999, 18, 785–789.

(34) Eckert-Maksic, M.; Glasovac, Z.; Troselj, P.; Kutt, A.; Rodima, T.; Koppel, I.; Koppel, I. A. Basicity of guanidines with heteroalkyl side chains in acetonitrile. Eur. J. Org. Chem. 2008, 2008, 5176–5184.

(35) Kubik, S.; Mungalpara, D. In Comprehensive Supramolecular Chemistry II ; Atwood, J. L., Ed.; Elsevier: Oxford, 2017; pp 293–310.

(36) Miyazaki, T.; Uchida, S.; Hatano, H.; Miyahara, Y.; Matsumoto, A.; Cabral, H. Guanidine-phosphate interactions stabilize polyion complex micelles based on flexible catiomers to improve mRNA delivery. Eur. Polym. J. 2020, 140, 110028.

(37) Kim, S.-H.; Semenya, D.; Castagnolo, D. Antimicrobial drugs bearing guanidine moieties: A review. Eur. J. Med. Chem. 2021, 216.

(38) Katayama, N.; Fukusumi, S.; Funabashi, Y.; Iwahi, T.; Ono, H. TAN-1057 A-D, new antibiotics with potent antibacterial activity against methicillin-resistant Staphylococcus aureus. Taxonomy, fermentation and biological activity. J. Antibiot. 1993, 46, 606–613.

(39) Heys, L.; Moore, C. G.; Murphy, P. J. The guanidine metabolites of and related compounds; isolation and synthesis. Chem. Soc. Rev. 2000, 29, 57–67.

(40) Belov, D. S.; Curreli, F.; Kurkin, A. V.; Altieri, A.; Debnath, A. K. Guanidine-containing phenyl-pyrrole compounds as probes for generating HIV entry inhibitors targeted to gp120. ChemistrySelect 2018, 3, 6450–6453.

(41) Cho, S. M.; Lee, H. K.; Liu, Q.; Wang, M.-W.; Kwon, H. J. A guanidine-based synthetic compound suppresses angiogenesis via inhibition of acid ceramidase. ACS Chem. Biol. 2019, 14, 11–19.

(42) Salvino, J. M.; Srikanth, Y. V. V.; Lou, R.; Oyer, H. M.; Chen, N.; Kim, F. J. Novel small molecule guanidine Sigma1 inhibitors for advanced prostate cancer. Bioorg. Med. Chem. Lett. 2017, 27, 2216–2220.

(43) Wu, Y.; Bi, Y.; Vavricka, C. J.; Sun, X.; Zhang, Y.; Gao, F.; Zhao, M.; Xiao, H.; Qin, C.; He, J.; Liu, W.; Yan, J.; Qi, J.; Gao, G. F. Characterization of two distinct neuraminidases from avian-origin human-infecting H7N9 influenza viruses. Cell Res. 2013, 23, 1347–1355.

(44) Artymiuk, P. J.; Blake, C. C. F.; Rice, D. W.; Wilson, K. S. The structures of the monoclinic and orthorhombic forms of hen egg-white lysozyme at 6 Å resolution. Acta Crystallogr. Sect. B 1982, 38, 778–783.

(45) Huang, J.; Rauscher, S.; Nawrocki, G.; Ran, T.; Feig, M.; de Groot, B.; Grubmueller, H.; MacKerell, A., Jr. CHARMM36m: an improved force field for folded and intrinsically disordered proteins. Nat. Methods 2017, 14, 71–73.

(46) Vanommeslaeghe, K.; Hatcher, E.; Acharya, C.; Kundu, S.; Zhong, S.; Shim, J.; Darian, E.; Guvench, O.; Lopes, P.; Vorobyov, I.; MacKerell, A. D., Jr. CHARMM general force field: a force field for drug-like molecules compatible with the CHARMM all-atom additive biological force fields. J. Comput. Chem. 2010, 31, 671–690.

(47) Jorgensen, W.; Chandrasekhar, J.; Madura, J.; Impey, R.; Klien, M. Comparison of simple potential functions for simulating liquid water. J. Chem. Phys. 1983, 79, 926–935.

(48) Phillips, J.; Braun, R.; Wang, W.; Gumbart, J.; Tajkhorshid, E.; Villa, E.; Chipot, C.; Skeel, R.; Kale, L.; Schulten, K. Scalable molecular dynamics with NAMD. J. Comput. Chem. 2005, 26, 1781–1802.

(49) Darden, T.; York, D.; Pedersen, L. Particle Mesh Ewald - an N log(N) method for Ewald sums in large systems. J. Chem. Phys. 1993, 98.

(50) Berendsen, H.; van der Spoel, D.; van Drunen, R. GROMACS: A message-passing parallel molecular dynamics implementation. Comput. Phys. Commun. 1995, 91, 43–56.

(51) Murtola, T.; Schulz, R.; Páll, S.; Smith, J. C.; Hess, B.; Lindahl, E. GROMACS: High performance molecular simulations through multi-level parallelism from laptops to supercomputers. SoftwareX 2015, 1–2, 19–25.

(52) Bussi, G.; Donadio, D.; Parrinello, M. Canonical sampling through velocity rescaling. J. Chem. Phys. 2007, 126.

(53) Parrinello, M.; Rahman, A. Polymorphic transitions in single-crystals - a new molecular-dynamics method. J. Appl. Phys. 1981, 52, 7182–7190.

(54) Lindorff-Larsen, K.; Piana, S.; Palmo, K.; Maragakis, P.; Klepeis, J. L.; Dror, R. O.; shaw, D. E. Improved side-chain torsion potentials for the Amber ff99SB protein force field. Proteins 2010, 78, 1950–1958.

(55) Wang, J.; Wolf, R.; Caldwell, J.; Kollman, P.; Case, D. Development and testing of a general amber force field. J. Comput. Chem. 2004, 25, 1157–1174.

(56) Wang, J.; Wang, W.; Kollman, P. A.; Case, D. A. Automatic atom type and bond type perception in molecular mechanical calculations. J. Mol. Graph. 2006, 25, 247–260.

(57) Case, D.; Cheatham, T.; Darden, T.; Gohlke, H.; Luo, R.; Merz, K.; Onufriev, A.; Simmerling, C.; Wang, B.; Woods, R. The Amber biomolecular simulation programs. J. Comput. Chem. 2005, 26, 1668–1688.

(58) Davies, R.; Neuberger, A.; Wilson, B. The dependence of lysozyme activity on pH and ionic strength. Biochim. Biophys. Acta - Enzymology 1969, 178, 294–305.

(59) Held, J.; van Smaalen, S. The active site of hen egg-white lysozyme: flexibility and chemical bonding. Acta Crystallogr. Sect. D-Struct. Biol. 2014, 70, 1136–1146.

(60) Bradley, J.; O’Meara, F.; Farrell, D.; Nielsen, J. E. Highly perturbed pK<sub>a</sub> values in the unfolded state of hen egg white lysozyme. Biophys. J. 2012, 102, 1636–1645.

(61) Bonomi, M. et al. Promoting transparency and reproducibility in enhanced molecular simulations. Nat. Methods 2019, 16, 670–673.

(62) Tribello, G. A.; Bonomi, M.; Branduardi, D.; Camilloni, C.; Bussi, G. PLUMED 2: New feathers for an old bird. Comput. Phys. Commun. 2014, 185, 604–613.

(63) Tang, Z.; Chang, C.-e. A. Binding thermodynamics and kinetics calculations using chemical host and guest: A comprehensive picture of molecular recognition. J. Chem. Theory Comput. 2018, 14, 303–318.

(64) Hänggi, P.; Talkner, P.; Borkovec, M. Reaction-rate theory - 50 years after Kramers. Rev. Mod. Phys. 1990, 62, 251–341.

(65) Scherer, M. K.; Trendelkamp-Schroer, B.; Paul, F.; Pérez-Hernández, G.; Hoffmann, M.; Plattner, N.; Wehmeyer, C.; Prinz, J.-H.; Noé, F. PyEMMA 2: A software package for estimation, validation, and analysis of Markov models. J. Chem. Theory Comput. 2015, 11, 5525–5542.

(66) Metzner, P.; Schuette, C.; Vanden-Eijnden, E. Transition path theory for Markov jump processes. Multiscale Model. Simul. 2009, 7, 1192–1219.

(67) Noe, F.; Schuette, C.; Vanden-Eijnden, E.; Reich, L.; Weikl, T. R. Constructing the equilibrium ensemble of folding pathways from short off-equilibrium simulations. Proc. Natl. Acad. Sci. U. S. A. 2009, 106, 19011–19016.

(68) Copeland, R. A.; Pompliano, D. L.; Meek, T. D. Opinion - Drug-target residence time and its implications for lead optimization. Nat. Rev. Drug Discov. 2006, 5, 730–739.

(69) Copeland, R. A. The drug-target residence time model: a 10-year retrospective. Nat. Rev. Drug Discov. 2016, 15, 87–95.

(70) Schuetz, D. A. et al. Kinetics for Drug Discovery: an industry-driven effort to target drug residence time. Drug Discov. Today 2017, 22, 896–911.

