## Supplementary Figures for "pH Effect on Ligand Binding to an Enzyme Active Site"

### Supplementary Information

Table S1: Non-bonded $Gdm^+$ parameters used in the AMBER simulations

| Atom | $\sigma$ (Å) | $\epsilon$ (kJ/mol) | Charge ($e_0$) |
|---|---|---|---|
| C | 0.3399 | 0.3598 | 1.0839 |
| N | 0.3249 | 0.7112 | -1.0137 |
| H | 0.1069 | 0.0656 | 0.4928 |

Table S2: Average mean first passage time (MFPT) for $Gdm^+$-protein complexes obtained from the Markov state model (MSM)

| | **U** | **A** | **PB** | **B1** | **B2** | **D** | **T** |
|---|---|---|---|---|---|---|---|
| **U** | 0.00 | 32.26 | 13.64 | 4.49 | 4.35 | 11.13 | 375.06 |
| **A** | 41.20 | 0.00 | 8.67 | 4.83 | 4.35 | 11.82 | 379.44 |
| **PB** | 43.30 | 31.61 | 0.00 | 3.38 | 3.86 | 14.64 | 380.82 |
| **B1** | 43.58 | 34.76 | 68.69 | 0.00 | 4.87 | 12.32 | 378.62 |
| **B2** | 43.12 | 31.84 | 84.75 | 5.27 | 0.00 | 12.82 | 373.41 |
| **D** | 45.33 | 35.04 | 92.07 | 6.42 | 6.32 | 0.00 | 370.60 |
| **T** | 54.73 | 46.20 | 105.26 | 16.03 | 8.21 | 17.17 | 0.00 |

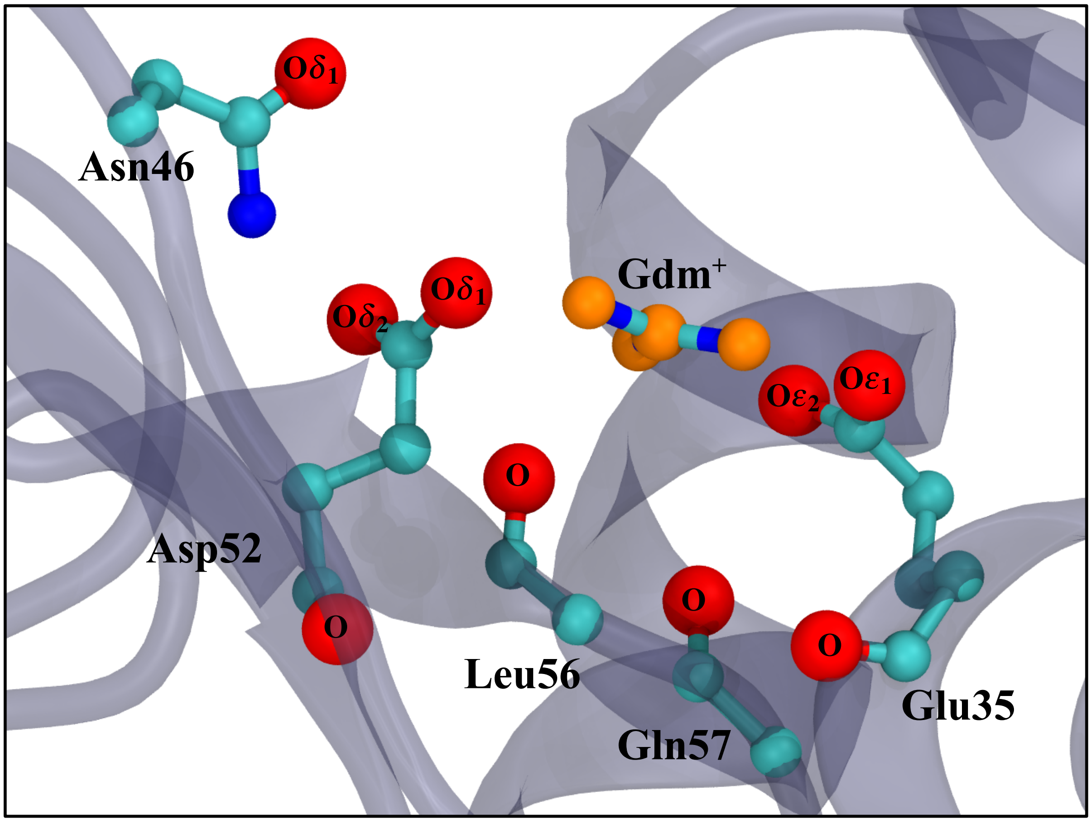


Figure S1: Residues in the vicinity of the binding pocket of HEWL. Distances between the oxygen atoms (marked in red) and the heavy atoms of $Gdm^+$ (orange) were selected as the input variable for constructing the MSM.

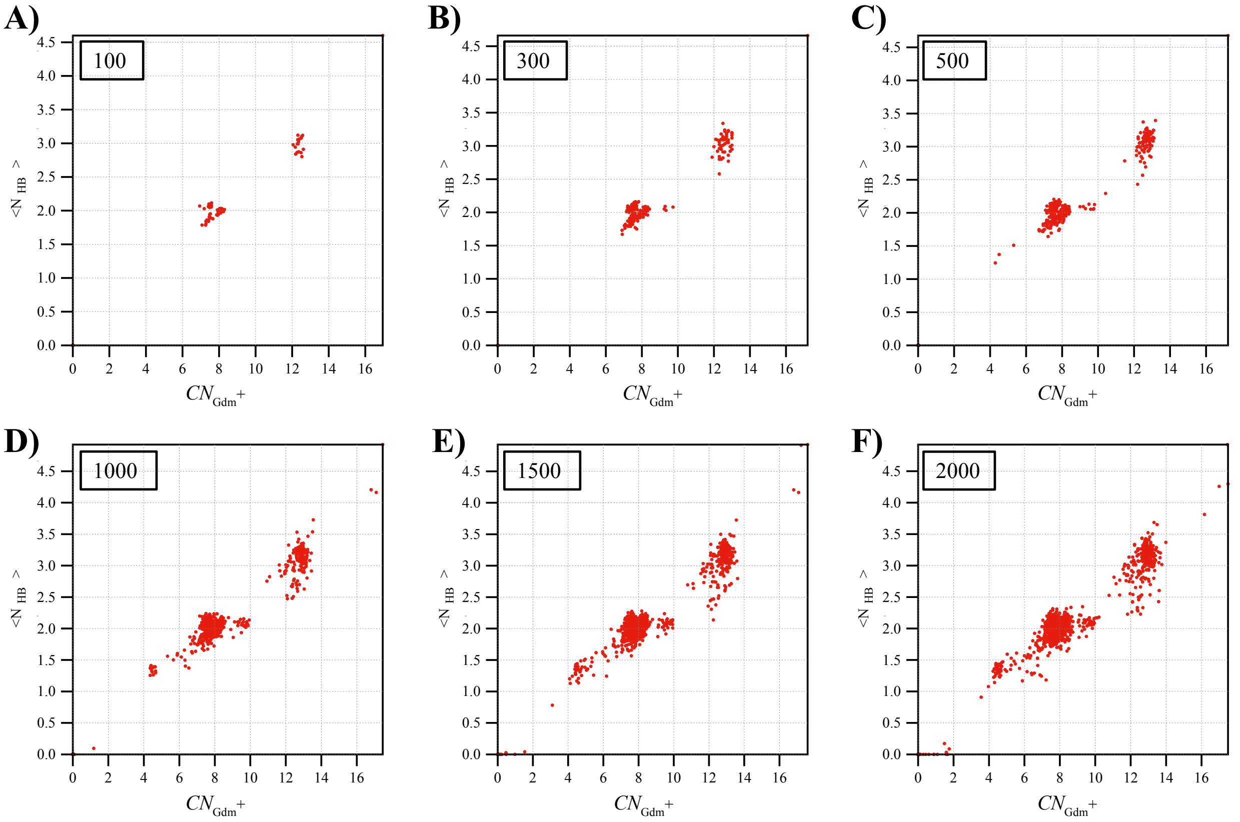


Figure S2: Projection of clusters onto $\langle N_{HB} \rangle$ and $CN_{Gdm^+}$.

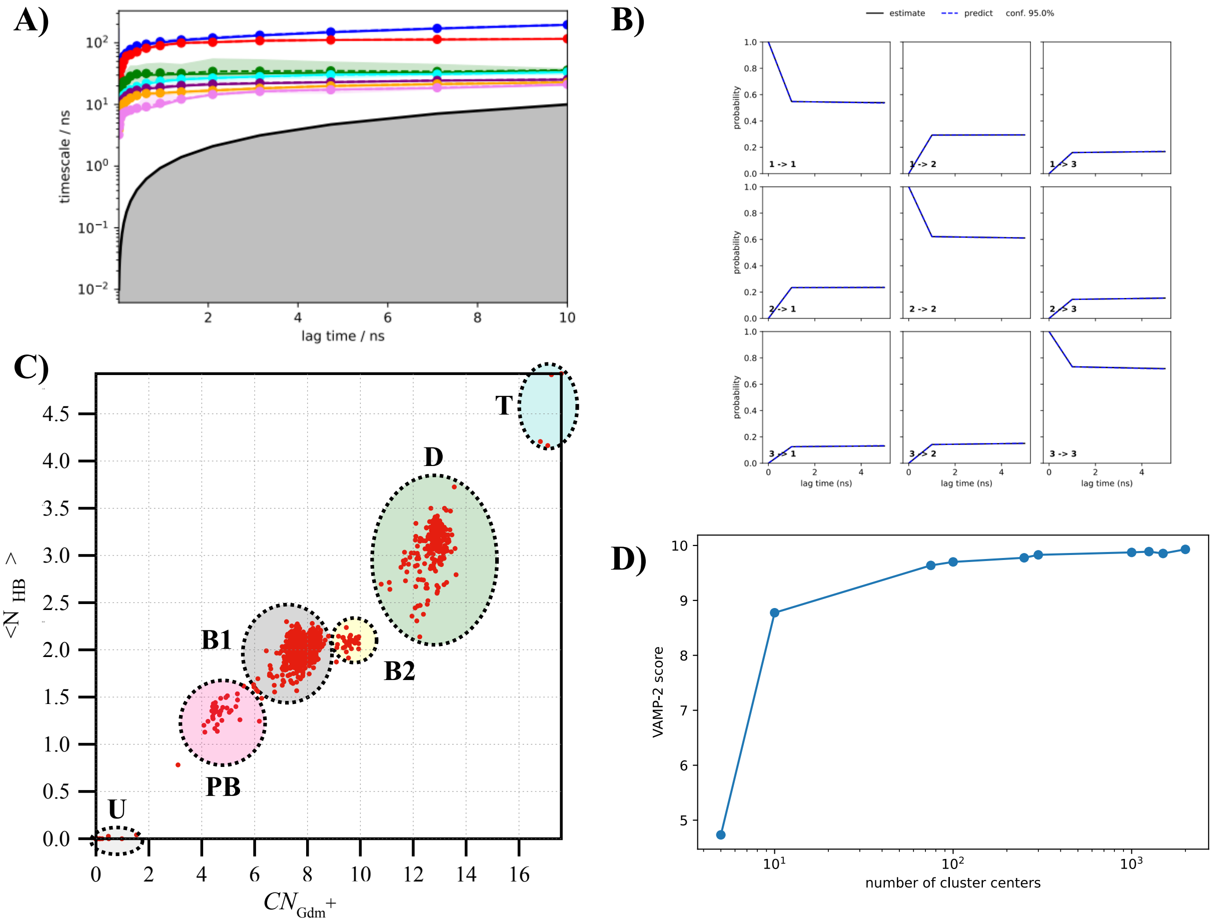


Figure S3: (A) The implied timescales plot showing the seven slowest processes up to a lag time of 10 ns. The separation between the first two and the other five timescales shows that MSM resolves the three state processes. The timescales level off at around 1 ns, indicating that 1 ns is the appropriate lag time to construct the MSM. (B) The Chapman-Kolmogorov (CK) test to validate the MSM. For the three state processes, the CK test shows that the MSM is valid at all lag times up to 5 ns. (C) Coarse-grained assignment of clusters to the various bound states of the $Gdm^+$-HEWL complex. (D) VAMP score as a function of cluster size.

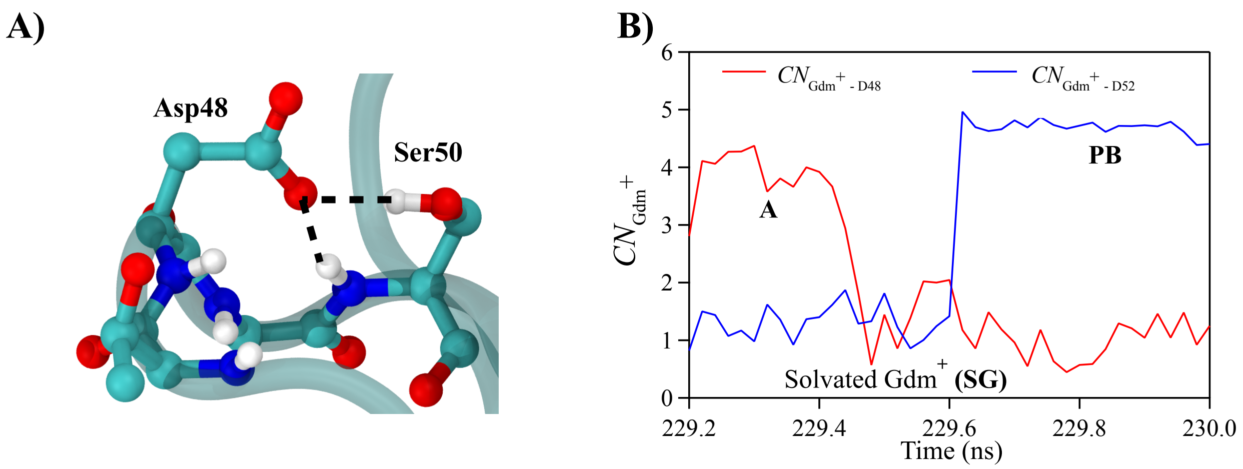


Figure S4: (A) The hydrogen bond network between Asp48 and Ser50. (B) The coordination number ($CN_{\mathrm{Gdm}^+}$) between a $\mathrm{Gdm}^+$ and residues Asp48 and Asp52 in states **A** (red) and **PB** (blue), respectively. The dip in $CN_{\mathrm{Gdm}^+}$ indicates that during the transition of a $\mathrm{Gdm}^+$ between states **A** and **PB**, the $\mathrm{Gdm}^+$ first moves away from Asp48 and is completely solvated before forming hydrogen-bonded interactions with Asp52.

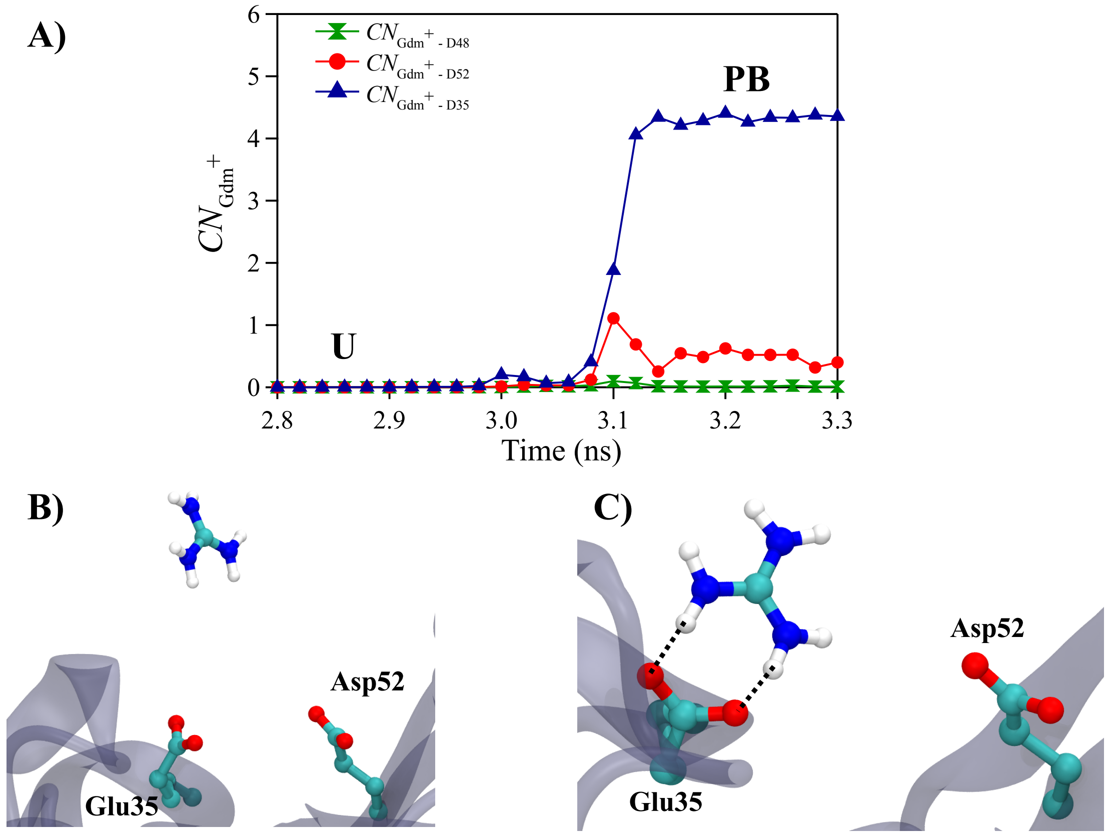


Figure S5: Direct transitions from state **U** to **PB** were observed in $Gdm^+$ binding to the HEWL active site. (A) Coordination number ($CN_{\text{Gdm}^+}$) between a $Gdm^+$ and Asp48 (green hourglass), Asp52 (red circles), and Glu35 (blue triangles), respectively. The $CN_{\text{Glu35-Gdm}^+}$ shows a sudden increase, while $CN_{\text{Asp48-Gdm}^+}$ and $CN_{\text{Asp52-Gdm}^+}$ remain low, indicating $Gdm^+$ binds directly to Glu35 from solution. (B) Representative configurations where $Gdm^+$ from solution binds to Glu35.

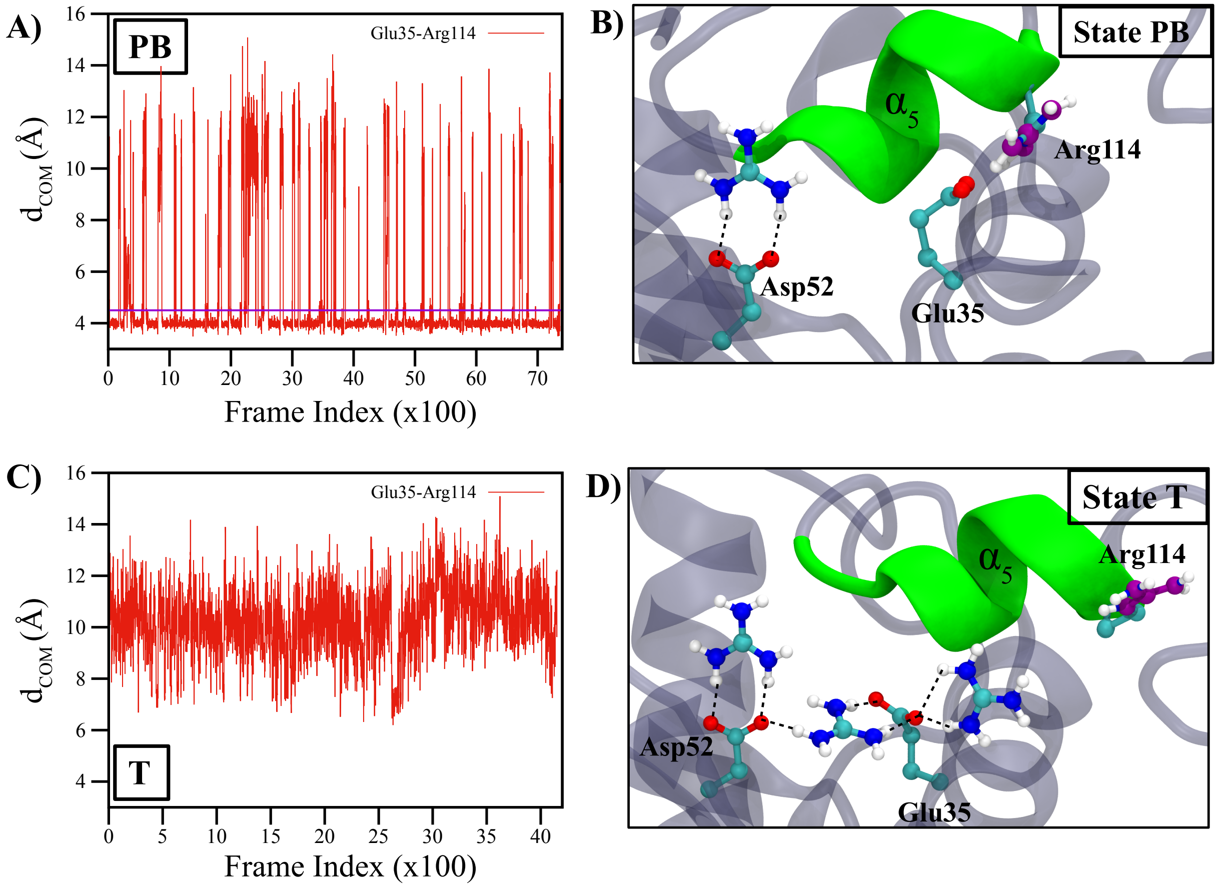


Figure S6: (A) and (C) Distance between the center of mass of Glu35 H-bond acceptor and center of mass of Arg114 H-bond donor in states A and T, respectively. Distances below a cutoff of 4.5 Å (magenta) indicate the formation of a hydrogen bond. (B) and (D) Representative configurations corresponding to the states A and T. The hydrogen bond distances between the carboxylic groups and the $Gdm^+$ ions are marked with black dashes. The guanidinium group of Arg114 has been marked in purple to distinguish it from $Gdm^+$ ions.

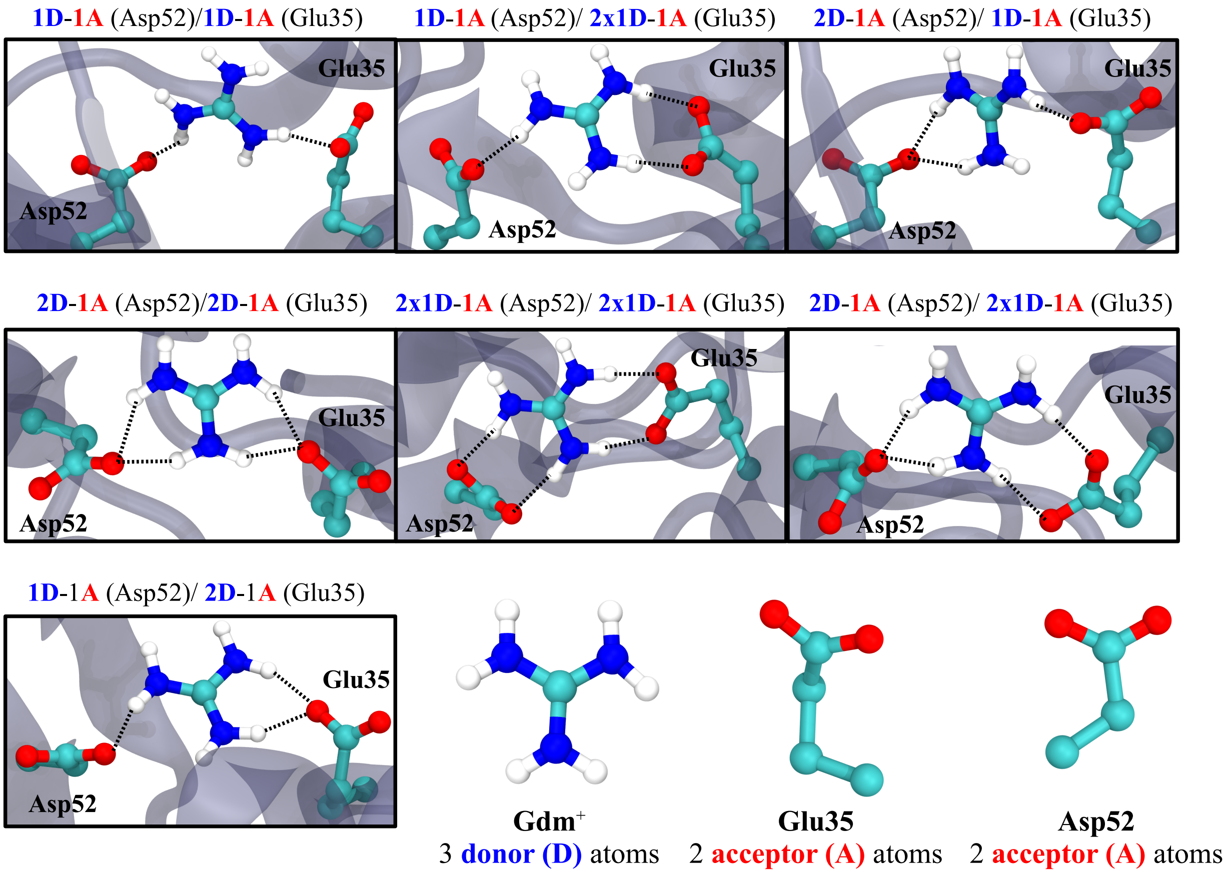


Figure S7: Various configurations of hydrogen bonds present between the active site of HEWL and a $Gdm^+$ in either state **B1** or **B2**.

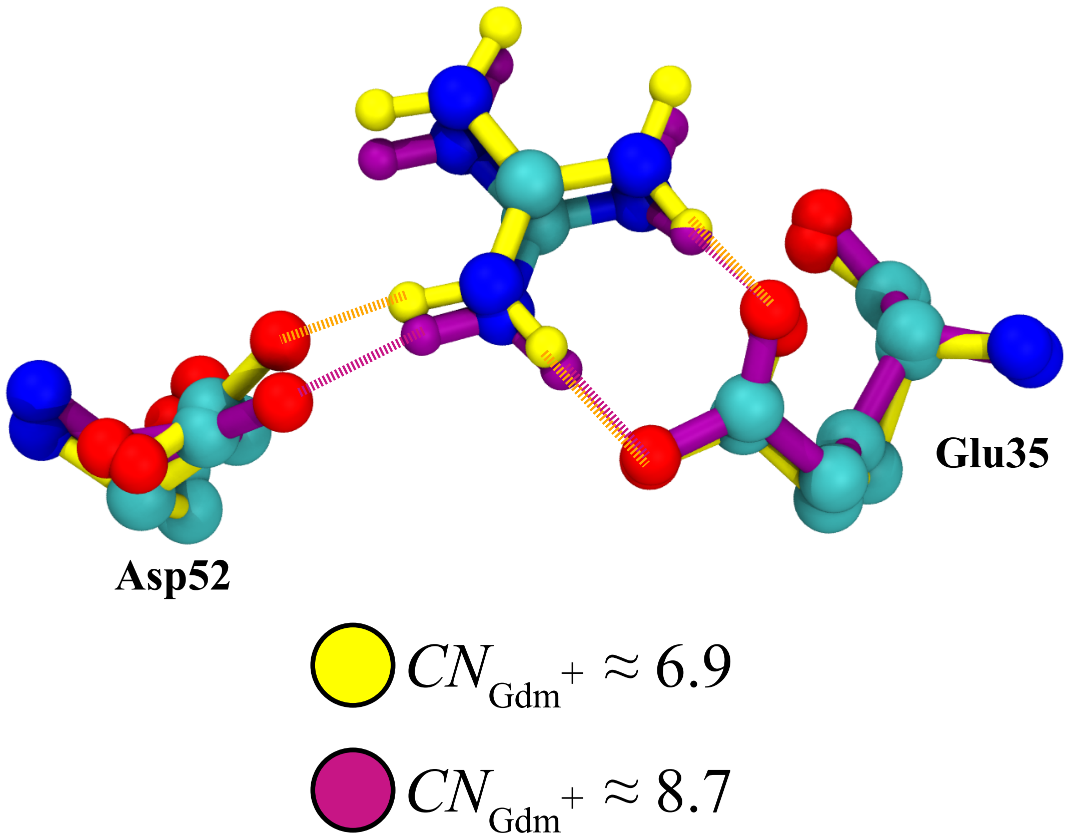


Figure S8: Superposition of the active site-$Gdm^+$ complex in states **B1** and state **B2** are shown with yellow and purple bonds, respectively. A small change in orientation of the carboxylate group of resid Asp52 results in $CN_{\mathrm{Gdm}^+}$ from 6.9 to 8.7.

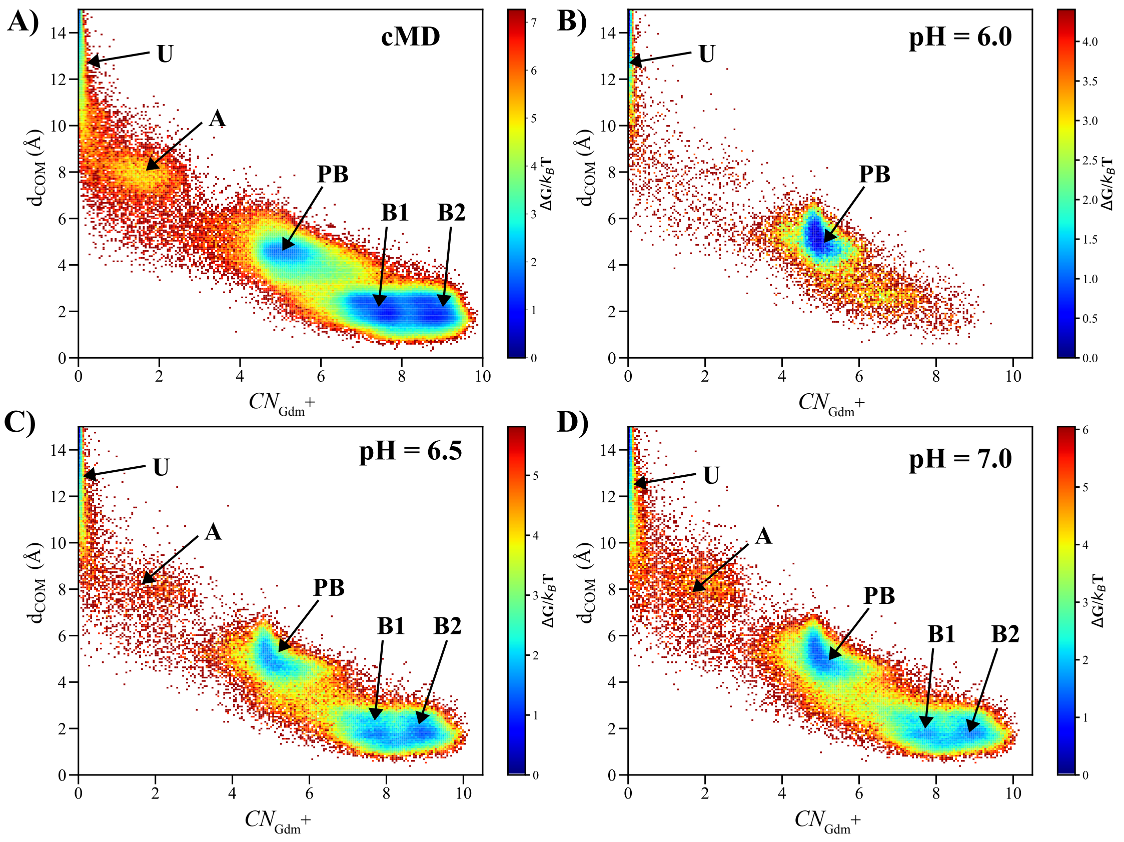


Figure S9: (A) The 2D FES projected onto $\mathrm{d_{COM}}$ and $CN_{\mathrm{Gdm}^+}$ for low [$\mathrm{Gdm}^+$]. The minima corresponding to the various configurations of $\mathrm{Gdm}^+$ are marked on the free energy surface. The minima occur in the same locations observed in the FES obtained using high [$\mathrm{Gdm}^+$]. However, at low pH (= 6.0), the basin corresponding to state **A** is missing, while shallower basins indicate lower binding probability at higher pH.

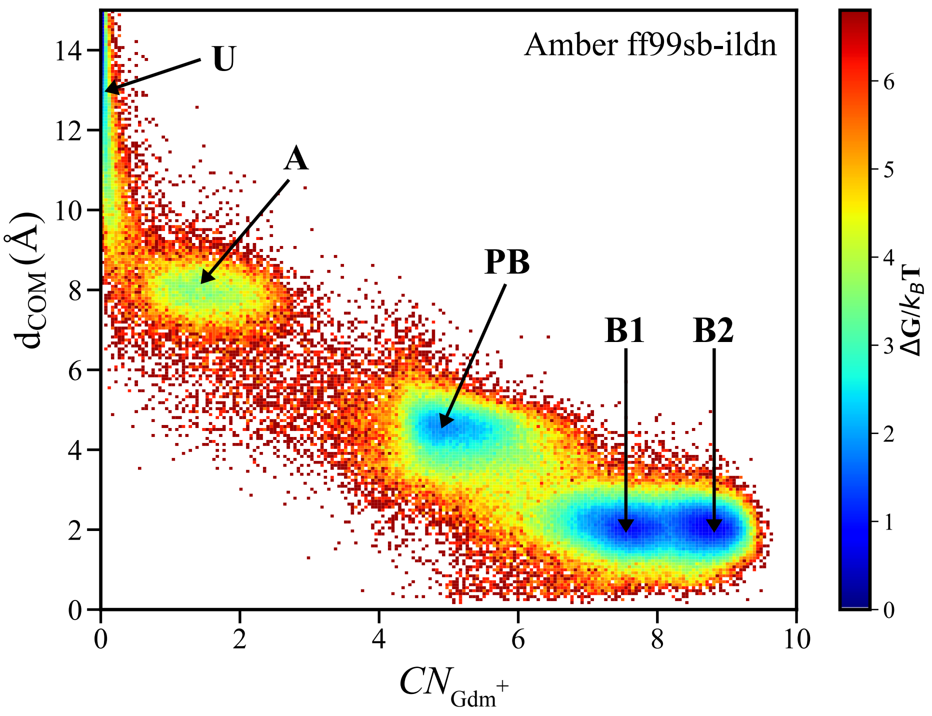


Figure S10: (A) The 2D FES obtained using the Amber ff99sb-ildn for the protein and GAFF for $Gdm^+$. The FES is projected onto $d_{COM}$ and $CN_{Gdm^+}$. The minima corresponding to the various configurations of $Gdm^+$ are marked on the FES.

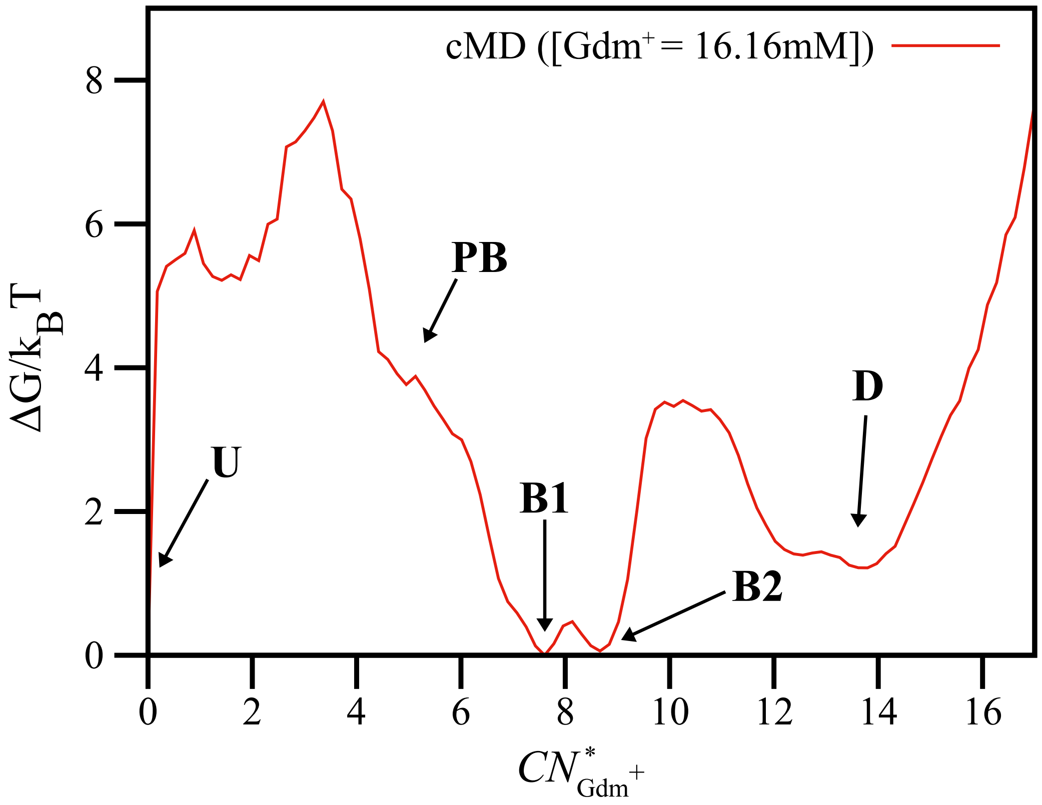


Figure S11: (A) The FES obtained using a low concentration of $Gdm^+$, $[Gdm^+]$ = 16 mM. The FES is projected onto $CN^*_{\mathrm{Gdm}^+}$. The minima corresponding to the various configurations of $Gdm^+$ are marked on the FES.

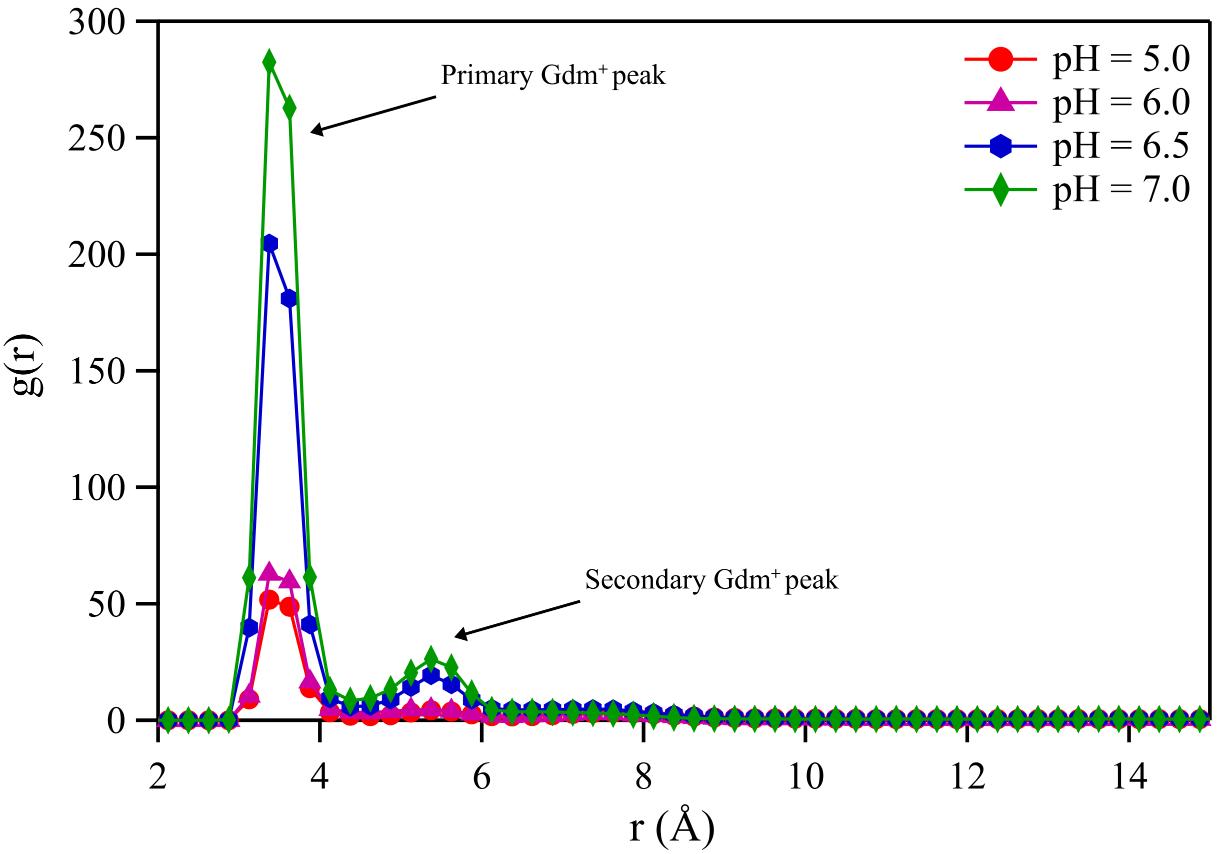


Figure S12: The radial distribution function, $g(r)$, between the center of mass of $Gdm^+$ and the carboxylate groups present in the HEWL active site is plotted for different pH values. At lower pH, the peak height at $\approx$ 3 Å (corresponding to a single $Gdm^+$ in the active site vicinity) is small, indicating that the binding of $Gdm^+$ to the protein active site is infrequent due to the high protonation rate of Glu35. At higher pH values ($>$ 6.5), the height of the primary peak increased, indicating stable binding. The secondary peak around 5 Å suggests another $Gdm^+$ is present at the binding site.

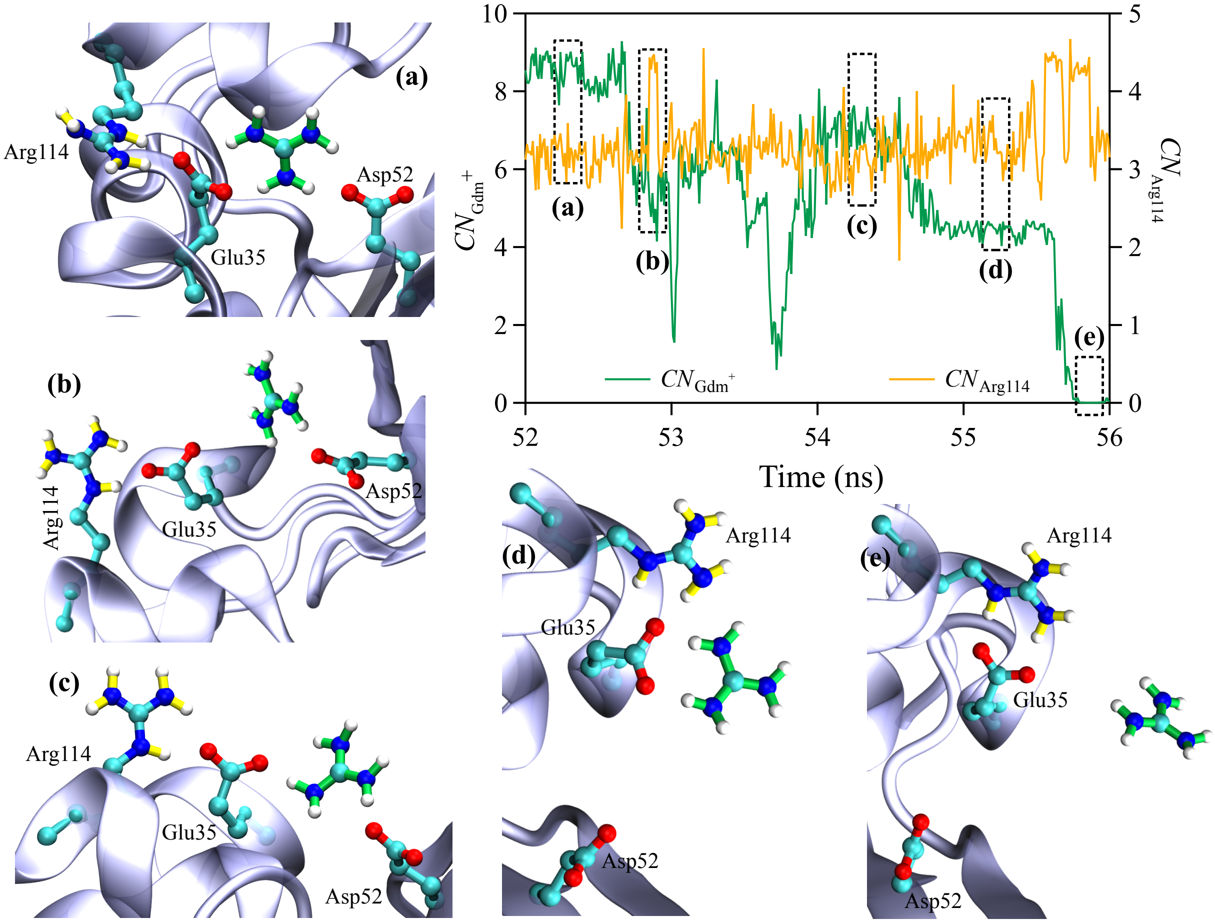


Figure S13: Secondary dissociation pathway of $Gdm^+$. The guanidine group of Arg114 (yellow) destabilizes the binding competent active site conformation by twisting the carboxylate group of Glu35 away from the active site as evidenced by the increase in $CN_{\text{Arg114}}$ (yellow). This weakens the bridging hydrogen-bonded interactions between $Gdm^+$ and the active site residues seen from the corresponding decrease in $CN_{\text{Gdm}^+}$ (green). The insets depict representative structures of key transitions in the unbinding pathway.
